# High-throughput Extracellular Matrix Proteomics of Human Lungs Enabled by Photocleavable Surfactant and diaPASEF

**DOI:** 10.1101/2023.08.17.553758

**Authors:** Elizabeth F. Bayne, Kevin M. Buck, Anna G. Towler, Yanlong Zhu, Melissa Pergande, Tianhua Zhou, Scott Price, Kalina J. Rossler, Vanessa Morales-Tirado, Sarah Lloyd, Fei Wang, Yupeng He, Yu Tian, Ying Ge

## Abstract

The extracellular matrix (ECM) is a complex assembly of proteins that provide interstitial scaffolding and elastic recoil to human lungs. The pulmonary extracellular matrix (ECM) is increasingly recognized as an independent bioactive entity by creating biochemical and mechanical signals that influence disease pathogenesis, making it an attractive therapeutic target. However, the pulmonary ECM proteome (“matrisome”) remains challenging to analyze by mass spectrometry due to its inherent biophysical properties and relatively low abundance. Here, we introduce a strategy designed for rapid and efficient characterization of the human pulmonary ECM using the photocleavable surfactant Azo. We coupled this approach with trapped ion mobility MS with diaPASEF to maximize depth of matrisome coverage. Using this strategy, we identify nearly 400 unique matrisome proteins with excellent reproducibility that are known to be important in lung biology, including key insoluble ECM proteins.

## Introduction

The extracellular matrix (ECM) is a complex assembly of proteins that provide interstitial scaffolding and elastic recoil to the lungs.^1^ The pulmonary ECM is increasingly recognized as an independent bioactive entity that provides biochemical and mechanical signals that influence disease pathogenesis,^2^ making it an attractive therapeutic target.^3, 4^ Dysregulation of the pulmonary ECM has been linked to fibrotic disease such as idiopathic pulmonary fibrosis (IPF) and lung cancer.^4, 5^ Efforts to characterize the extracellular matrix proteome have resulted in the generation of an *in silico* predictive network called the “matrisome,” including genes encoding structural proteins classified as “core matrisome,” and those involved in ECM regulation and signaling known as “matrisome associated” proteins.^6–8^ The database is composed of 274 core matrisome proteins, including collagens, glycoproteins, and proteoglycans, and 753 matrisome associated proteins, including ECM regulators, secreted factors, and ECM-affiliated proteins.^8–10^

Analysis of the pulmonary matrisome remains challenging due to its insoluble nature and relatively low abundance within the proteome.^7, 11^ Many key structural components of the ECM, such as elastin and collagen fibers, are minimally soluble due to the properties of their macromolecular assembly including dense networks of covalently cross-linked high molecular weight proteins.^12, 13^ The ECM is highly post-translationally modified, especially covalently-linked glycans and hydroxylated residues, which further limits protein solubility and digestion efficacy.^14^ Additionally, many matrisome proteins are lower abundance compared to other proteins within tissues, such as cytosolic and nuclear proteins.^11, 15^ Previous protocols have addressed these issues by carrying out lengthy chemical digestion steps to access the insoluble ECM^12, 16^ and using multiple fractions containing MS-incompatible reagents such as SDS and 8M urea,^17–19^ necessitating additional sample cleanup. These protocols are time-consuming in terms of lengthy chemical digestion steps, sample cleanup, and number of fractions for MS analysis, which may limit their application to large-scale experiments of clinical samples for biomarker discovery. Thus, we aim to develop a high-throughput (<1 day) proteomics workflow to reproducibly analyze the pulmonary matrisome from human lung tissue.

Here, we introduce a proteomics strategy to streamline the enrichment of matrisome proteins from the lungs using a photocleavable surfactant Azo^20^ and data-independent acquisition with parallel accumulation and serial fragmentation (diaPASEF)^21^ on a trapped ion mobility mass spectrometer. Azo enables rapid enzymatic digestion times (as fast as 30 min) and only 5 min irradiation with UV light to cleave the surfactant into MS-compatible degradation products.^22, 23^ We previously developed a MS-based method for analyzing the matrisome of mouse mammary tumor tissue using Azo, utilizing high pH reverse phase fractionation and data-dependent acquisition on a quadrupole time of flight MS.^24^ Importantly, Azo outperformed typical chaotropic extraction reagent 8M urea in solubilizing fibril-forming and basement membrane collagens.^24^ Hence, we utilized Azo in this study to enable efficient solubilization of the pulmonary ECM in a single fraction, eliminating the need for multiple MS-incompatible reagents such as cyanogen bromide and urea. To further reduce the number of fractions required to analyze the matrisome, the resulting peptides are analyzed by diaPASEF to address issues with dynamic range and sensitivity and capture the matrisome without extensive pre-fractionation.

Using this rapid ECM proteomics method, we identify nearly 400 unique matrisome proteins with excellent reproducibility known to be important in lung biology, including key insoluble ECM proteins. We demonstrate successful enrichment of collagens, proteoglycans, and glycoproteins using Azo, including the insoluble ECM (iECM). Overall, this approach enables reproducible and rapid access to the pulmonary ECM from human lungs. The single-step solubilization of the insoluble matrisome with Azo will enable understanding of the highly complex three-dimensional architecture of lung tissue. This discovery-based strategy can be used to generate hypotheses and identify potential therapeutic targets and advance understanding of ECM-mediated lung pathogenesis.

## Materials and Methods

### Reagents and chemicals

Unless otherwise noted, reagents and chemicals were purchased from Sigma-Aldrich, Inc. (St. Louis, MO, USA). HPLC-grade water and acetonitrile were purchased from Fisher Scientific (Fair Lawn, NJ, USA)

### Lung Tissue Collection

Central and peripheral lung tissue from the left upper lobe of donors with no history of pulmonary disease were obtained from the University of Wisconsin Organ Procurement Organization (**Table S1** details clinical characteristics). Tissue collection procedures were approved by Institutional Review Board of the University of Wisconsin-Madison (Study # 2011-0868). Lung tissue was dissected into peripheral and central tissues of each lobe while fresh, immediately snap frozen in liquid nitrogen, and stored at -80 °C until analysis.

### Preparation of Lung Tissue

Approximately 30 mg of frozen lung tissue was weighed into a 1.5 mL tube at 4 °C. The tissue was cryopulverized (Cellcrusher Kit, ThermoFisher Scientific) using liquid nitrogen and placed into 1.5 mL centrifuge tubes. Dulbecco’s Phosphate Buffered Saline (0.9% NaCl) with the addition of 1x HALT Protease and Phosphatase inhibitor was added to the cryopulverized tissue. The samples were incubated for 5 min on a nutator at 4 °C. The saline-tissue mixture was centrifuged for 1 min at 1,000 × *g* to gently pellet the tissue. The supernatant was removed and discarded. The rinse was repeated with 300 µL saline mixture.

### Extraction of the Lung ECM Proteome

Proteins from the pelleted lung tissue were extracted using a buffer containing 40 mM HEPES (pH 7.9), 60 mM NaF, 1x HALT Protease and Phosphatase Inhibitor, 1 mM Na_3_VO_4_, 1 mM PMSF, 1 mM DTT, and 25 mM EDTA. Tissue was manually homogenized in buffer using a handheld Teflon homogenizer, centrifuged at 18,000 × *g* for 15 min (4 °C), and the supernatant was transferred to a separate tube. This process was repeated twice. The supernatant from the three rounds of cytosolic extraction was pooled to create the “Decell extract” and later processed for bottom-up proteomics analysis. The pooled extract was buffer exchanged into 25 mM Ammonium Bicarbonate (ABC) using 10 kDa MWCO filters (Pierce Protein Concentrators, ThermoFisher Scientific).

Proteins from the remaining pellet were solubilized in 100 uL buffer containing 0.5% w/v Azo, 25 mM ABC, 1x HALT, 1 mM PMSF, 1 mM Na_3_VO_4_, 2.5 mM EDTA, and 1 mM DTT. The pellet was homogenized into the buffer using a handheld Teflon homogenizer. The solutions were probe sonicated (20% amplitude, 3 cycles, 3 sec each), heated to 95 °C for 30 min, and sonicated in a water bath for 60 min. The samples were centrifuged for 10 min at 20,000 × *g* at 24 °C and the supernatant (“Azo extract”) was transferred to a clean tube.

The Decell and Azo extracts were normalized using Pierce Bradford Plus Protein Assay (ThermoFisher Scientific), reduced, alkylated, and digested with Trypsin Gold (Promega, Madison, WI). Azo extracts were irradiated by UV light at 305 nm for 5 min to degrade the photocleavable surfactant. Samples were offline desalted using Pierce C18 Spin Tips & Columns (ThermoFisher Scientific) and reconstituted in 0.1% formic acid in water. Peptide concentration was assessed using Nanodrop One prior to LC-MS injection.

### Data Acquisition

Nanoflow LC-MS analysis of tryptic peptides was performed using a timsTOF Pro (Bruker Daltonics, Bremen, Germany)^25^ coupled to a nanoElute nano-flow UHPLC system (Bruker Daltonics, Bremen, Germany) using a Captivespray nano-electrospray ion source. 200 ng peptides were loaded onto an IonOpticks capillary C18 column (25 cm x 75 μm i.d., 1.7 μm) and peptides were separated at 55 °C using a 400 nL/min flow rate and a stepwise gradient of 2 to 17% B over 60 minutes and 17 to 25% B from 60 to 90 min, followed by a 30 min wash step at 85% B, totaling 120 min per run. The mass range for MS and MS/MS scans was 100-1700 *m/z*. The timsTOF Pro was operated in Parallel Accumulation and Serial Fragmentation (PASEF) mode^25^ and Data Independent Acquisition (DIA) mode (diaPASEF).^21, 26^ Ion mobility resolution was set to 0.60-1.60 V*s/cm. Collisional induced dissociation (CID) energies ranged from 20-59 eV and scaled on ion mobilities, 1/k0. Peptide loading was assessed by comparing the intensity of the total ion current to 200 ng of a human K562 whole cell lysate standard (Promega, Madison, Wisconsin, USA) prepared at 0.2 μg/μL.

### Data Analysis

Tandem mass spectra were searched using a library-free approach by DIA-NN v1.8.1.^27^ A spectral library was generated using a FASTA digest from the UniProt human database UP000005640 (canonical and isoform, accessed 6 October 2022). Peptides had a minimum of 7 amino acids with carbamidomethylation as a fixed modification and N-terminal acetylation and methionine oxidation as variable modifications. Enzyme specificity was set to trypsin/P and a maximum of 2 missed cleavages were allowed. The maximum mass tolerance for precursor ions was 20.0 ppm and fragment ions was 10.0 ppm. A precursor false discovery rate of 1% was used. Cross-run normalization was applied using Retention time-dependent mode.

Protein intensities were log_2_-transformed and proteins removed if not present in ≥ 2 of 3 extraction replicates. Proteins were matched by Official Gene Symbols to the Naba Matrisome Database.^10, 14^ BioVenn was used to determine Protein Group overlap.^28^ Protein sequence coverage was determined by the Protein Coverage Summarizer Tool from PNNL. Total Protein Analysis (TPA) was performed as previously described.^29^ Molecular weights for each protein were procured from UniProtDB. Pathway analysis and functional enrichment was performed on the dataset using the Database for Annotation, Visualization, and Integrated Discovery (DAVID).^30^ Gene ontology was performed by PANTHER^31^ Gene Ontology Cellular Component, and statistical overrepresentation of GOCC terms was performed by a Fisher’s Exact Test, and FDR-corrected *p* values were -log10x transformed and plotted. STRING analysis^32^ was performed on identified matrisome proteins.

## Results and Discussion

We have developed a rapid (<1 day) and facile method to reproducibly extract matrisome proteins directly from the human lung without extensive pre-fractionation and sample cleanup. We employed a single step solubilization of ECM proteins using the photocleavable surfactant Azo (4-hexylphenylazosulfonate).^22^ We coupled this approach with trapped ion mobility MS with diaPASEF to maximize depth of sample coverage, eliminating the need for sample pre-fractionation. Using this strategy, we identified nearly 400 unique matrisome proteins with high reproducibility, including key insoluble ECM proteins.

### The Enrichment of Extracellular Matrix Proteins from Human Lungs

Sample preparation includes the depletion of highly abundant blood and plasma proteins, fractionation of soluble proteins by decellularization, and enrichment of ECM from the remaining insoluble pellet using the photocleavable surfactant Azo (**Figure 1**). This extraction method yields two protein extracts analyzed by LC-MS/MS, referred to as the “Decell” and “Azo” extracts. The Decell extract is created by homogenization of lung tissue in a decellularization buffer to deplete soluble cytosolic and intracellular proteins. The Azo extract consists of proteins extracted from the residual insoluble pellet, which is immersed in Azo surfactant and subjected to sonication and heating to enable facile solubilization of ECM proteins. Importantly, a significant increase in the relative abundance of collagen species was achieved by Azo compared to traditional ECM extraction methods such as highly chaotropic 8 M urea.^24^ Additionally, Azo enables efficient sample throughput by requiring only 30 min for tryptic digestion and 5 minutes of irradiation by UV light to cleave into MS-compatible degradation products.^23^

**Figure 1.**
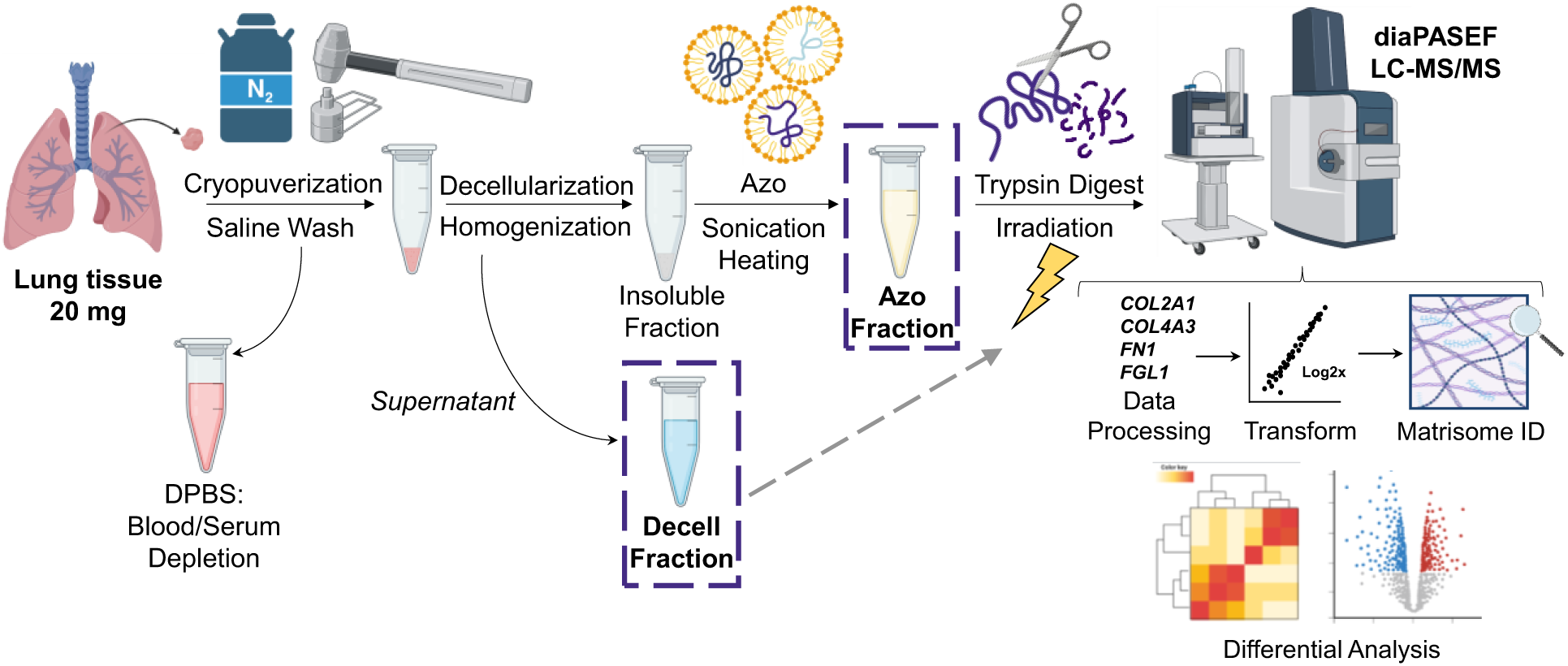
Experimental Workflow for the Enrichment of Extracellular Matrix Proteins from Human Lungs. Peripheral and central lung tissues of the left upper lobe were collected from adult donor lungs with no pulmonary history and snap frozen in liquid nitrogen. Tissue was cryopulverized, saline rinsed, and decellularized using pH-neutral buffer. Extracellular matrix proteins were enriched from the insoluble pellet using the anionic photocleavable surfactant Azo. Proteins were enzymatically digested and 200 ng tryptic peptides were analyzed by LC-MS/MS using Bruker timsTOF Pro with diaPASEF (parallel accumulation–serial fragmentation with data-independent acquisition). Tandem mass spectra were matched to an in-silico spectral library generated by the UniProt human database (DIA-NN v1.8.1 library-free mode, 1% FDR). Proteins were matched to the Naba MatrisomeDB. Data analysis and visualization was performed using BioVenn, Perseus, and R Studio.

To further enable deep sample coverage, we used the data independent acquisition with Parallel Accumulation and Serial Fragmentation (diaPASEF) mode on the timsTOF Pro.^21^ Data independent acquisition is a useful approach for analyzing the matrisome, since the matrisome sub-proteome is of relatively low abundance compared to intracellular proteins.^11^ As a result, we achieved deep proteome profiling routinely of human lung tissues, identifying approximately 6,000 protein groups from each fraction.

### Optimization of Sample Preparation

Highly abundant plasma proteins in lung tissue are present due to dense networks of capillaries surrounding the alveoli to perform gas exchange. Wide dynamic range of the blood proteome represents a hurdle for quantitation of lower-abundant lung and ECM proteins and may interfere with the quantitative ability of the MS method by dominating the mass spectral signal. To deplete highly abundant plasma proteins, cryopulverized lung tissue was repeatedly immersed in DPBS with protease and phosphatase inhibitors (**Figure S2A**). The tissue was gently pelleted by low rpm centrifugation (1,000 × *g*) and the DPBS was removed and discarded. We found this sequential wash depleted highly abundant plasma proteins such as human serum albumin (gene: *ALB*) effectively by SDS-PAGE (**Figure S2B**). LC-MS/MS samples prepared from the same lung tissue that was not washed prior to protein extraction compared to those that underwent the DPBS wash protocol showed an increase in overall Total Ion Chromatogram (TIC) intensity (**Figure S2C**). An average of 4,574 additional protein identifications were made in LC-MS/MS samples created from the washed tissue compared to the non-washed tissue (**Figure S2D**). Additionally, we found a significant reduction in the relative abundance of albumin in the washed samples (**Figure S2E, S2F**).

Following the removal of highly abundant plasma proteins, a decellularization buffer was added to extract cytosolic, nuclear, and ECM-affiliated proteins. The resulting protein extract, the “Decell extract,” was analyzed by LC-MS as part of our matrisome analysis. Previously, decellularization was performed using the surfactant Triton X-100.^24^ However, Triton is not MS-compatible, causes severe ionization suppression, and is difficult to remove prior to analysis.^33^ HEPES-based extraction buffer of pH 7.9 has been shown to successfully decellularize tissues^19^ and thus, we replaced Triton with a MS-compatible HEPES-based extraction buffer of pH 7.9. Upon comparison, Triton and HEPES showed similar performance for decellularization (**Figure S3**).

### Extracellular Matrix Protein Extraction is Highly Reproducible

Our method successfully enriches extracellular matrix proteins from human lungs with excellent reproducibility. In the Decell fraction, protein identifications ranged from 5,768-5,899 protein groups between three extraction replicates of lung tissue, corresponding to 5,518 proteins in common and 90% overlap (**Figure 2A**). Protein identifications in the Azo fraction ranged from 6,071-6,465 protein groups between three lung tissue extraction replicates, corresponding to 5,825 protein groups in common and an 87% overlap (**Figure 2B**). A total of 8,940 proteins were identified between the two fractions, including 2,031 proteins uniquely identified in the Decell extract and 2,680 proteins uniquely identified in the Azo extract (**Figure 2C**). Good spectral intensity and reproducibility was observed in the Decell fractions (**Figure S3A**) and Azo fractions (**Figure S3B**). Approximately 60,000 precursor ions were identified per run (**Figure S3C-D**) with normal, unimodal distribution of peptide intensities throughout the runs (**Figures S3E-F**). Strong correlation of protein intensity between all identified proteins in each fraction demonstrates strong extraction reproducibility. Pearson correlation coefficients between extractions ranged from 0.960-0.982, collectively (**Figures S3G, H**). As expected, the highest abundance proteins in the Decell fraction had overrepresented GOCC terms relating to intracellular space, including organelle identification and ribosomes (**Figure S4A, B**). The most abundant proteins in the Azo fraction corresponded to GOCC terms related to membrane and ECM proteins (**Figure S4C,D**).

**Figure 2.**
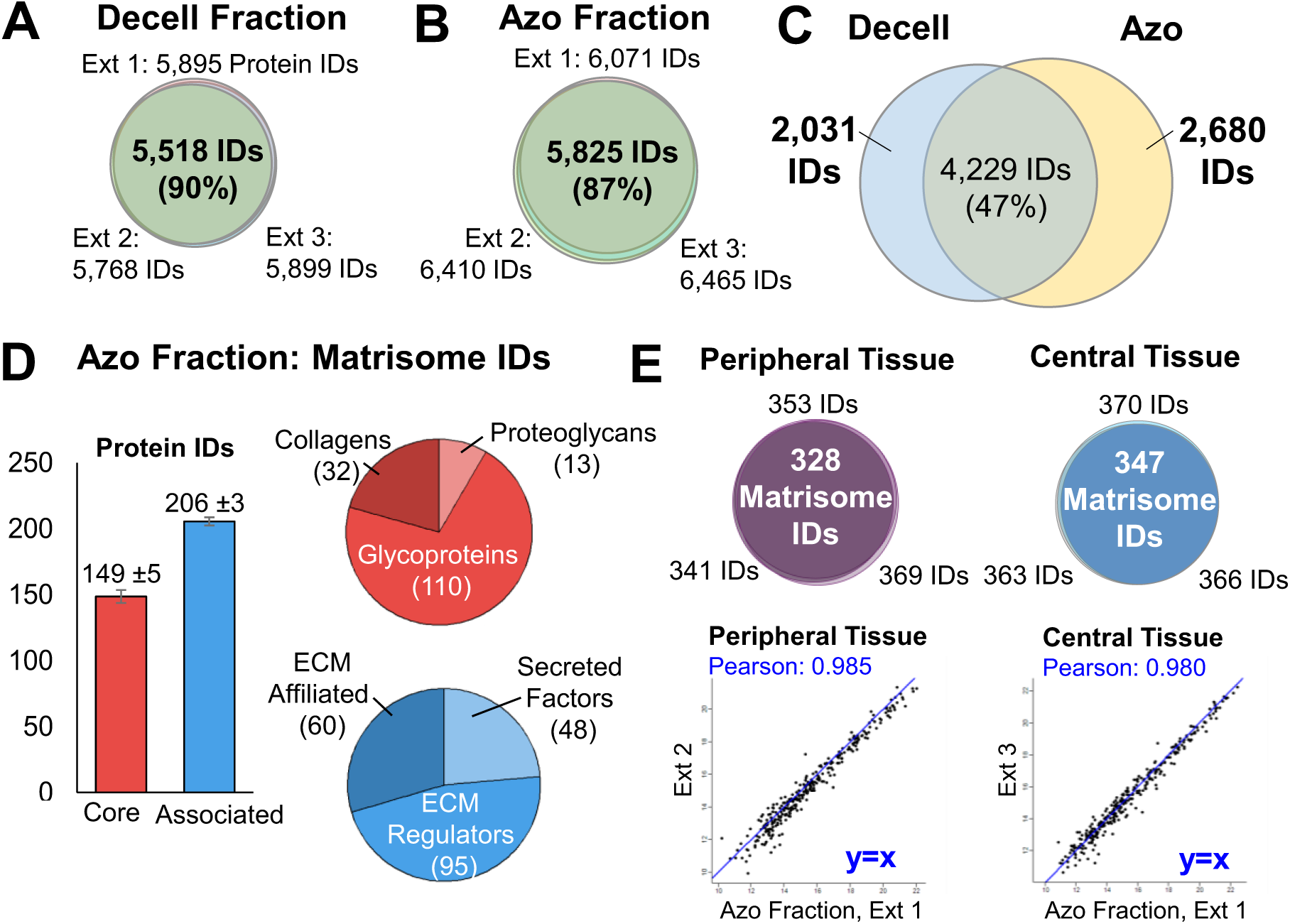
Extracellular Matrix Protocol is Highly Reproducible. **A)** Protein identifications in the Decell fraction ranged from 5,768-5,899 protein groups between three lung tissue extraction replicates, corresponding to 5,518 proteins in common and a 90% overlap. **B)** Protein identifications in the Azo fraction ranged from 6,071-6,465 protein groups between three lung tissue extraction replicates, corresponding to 5,825 protein groups in common and a 87% overlap. **C)** Comparison of protein identifications made in the Decell and Azo fractions. A total of 8,940 proteins were identified combining both fractions. 2,031 proteins were uniquely identified in the Decell fraction, and 2,680 proteins were uniquely identified in the Azo fraction. **D)** Matrisome protein identifications in the Azo Fraction. An average of 149 ±5 core matrisome proteins and 206 ±3 Matrisome-associated proteins were identified in the Azo fraction (Mean ±SE). The core matrisome protein identifications comprised 32 Collagen species, 13 Proteoglycans, and 110 Glycoproteins. The Matrisome-associated protein identifications included 60 ECM Affiliated proteins, 48 Secreted Factors, and 95 ECM Regulators. E) Matrisome protein identifications across (n=3) extraction replicates of peripheral and central lung tissues from the left upper lobe. In Peripheral tissues, 328 Matrisome proteins were identified in all 3 extraction replicates, and in central tissues, 347 matrisome proteins were shared between 3 replicates. F) log2x-transformed intensities of shared Matrisome proteins in each extraction replicate of Peripheral lung tissue (left), ranging from Pearson correlation coefficients ranging from 0.965-0.985 across replicates. In Central lung tissue (right), Pearson correlation coefficients ranged from 0.940-0.980 across replicates.

In the Azo fraction, we captured a total of 378 ECM proteins with high quantitative reproducibility (**Figure 2D**). An average of 149 ±5 core matrisome and 206 ±3 matrisome associated proteins were identified between extraction replicates (Mean ± SE). We identified 32 unique collagen species, 13 proteoglycans, 110 glycoproteins, 60 ECM-affiliated proteins, 48 secreted factors, and 95 ECM regulators in the Azo fraction. The majority of matrisome proteins identified in the Decell extract were matrisome-associated proteins, including ECM-affiliated proteins, ECM regulators, and secreted factors. In the Decell fraction, an average of 110 ±2 core matrisome proteins and 165 ±1 matrisome-associated proteins were identified between extraction replicates (**Figure S5**). The core matrisome species consisted of 15 collagens, 12 proteoglycans, and 89 glycoproteins. The matrisome associated proteins consisted of 53 ECM-affiliated proteins, 34 secreted factors, and 90 ECM regulators.

Our method demonstrated reproducible capture of matrisome proteins in both peripheral and central regions of the left upper lobe in human lungs, accounting for regional differences in ECM structures in the lung. In the Azo extract, 328 Matrisome proteins were identified from peripheral lung tissue (n=3 extraction replicates) and 347 matrisome proteins were recovered from central lung tissues (**Figure 2E**). Strong correlation of log2x-transformed protein intensities (0.965-0.985, Pearson) between extraction replicates for each anatomical region shows consistent recovery of the matrisome from the pellet (**Figure 2F**).

### Core Matrisome Proteins are Significantly Enriched in the Azo Fraction

Azo significantly enriched ECM proteins compared to the decellularized fraction. A total of 169 Core Matrisome Proteins were collectively identified in the Decell and Azo extracts (**Figure 3A**). Four proteins were identified exclusively in the Decell fraction, three of which belong to the insulin-like growth factor-binding protein family (genes: *IGFBP2*, *IGFBP3*, and *IGFBP6*). In the Azo fraction, 53 proteins, including alternatively-spliced isoforms, were uniquely identified including 16 collagen species, 34 glycoproteins, and 3 proteoglycans (summarized in **Table S3**). Notably, elastin alternatively-spliced isoforms 1, 5, and 11 were detected in this fraction. Alternatively spliced isoforms of elastin are thought to be associated with development and aging.^34^ Importantly, we detected 17 collagen species and 32 unique collagen isoforms, including prominent collagen species such as Collagen 6 (**Table S4**). In comparison to the predicted *in silico* Matrisome database,^35^ we identified 32 of 44 predicted collagen isoforms, 13 of 35 known proteoglycans, and 110 of 195 predicted glycoproteins from a single fraction (**Figure S6A**). This coverage is comparable to recent reports of multi-step extraction protocols from leading ECM groups.^18^ Additionally, our Azo-enabled matrisome coverage expands beyond experimentally observed core matrisome entries from human lung tissue in the Matrisome DB,^10^ with 61 unique core matrisome proteins that were not reported under human lung tissue (data sourced from Lansky *et. al.,* 2019,^36^ **Figure S6B**). Thus, our method provides a streamlined lung ECM proteomics approach and greatly expand the experimental matrisome knowledge database of human lung tissue.

**Figure 3.**
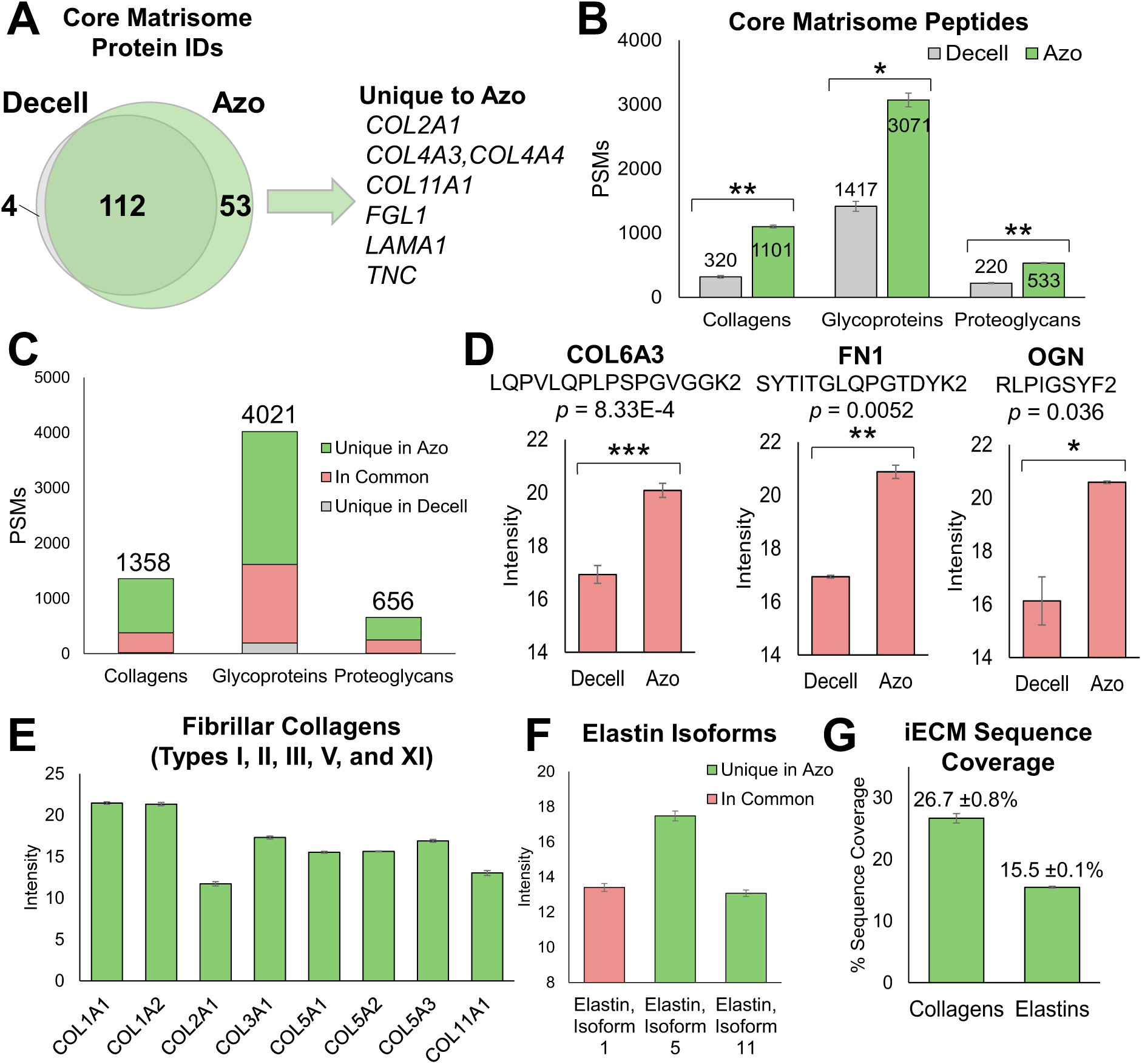
Core Matrisome Proteins are Significantly Enriched in the Azo Fraction. **A)** Core Matrisome protein identifications in the Decell and Azo fractions. Four unique protein IDs were found in the Decell fraction and 53 unique isoforms (including alternatively spliced isoforms) were found in the Azo fraction, for a total of 40 unique gene families. Representative gene families found unique to the Azo fraction are highlighted. **B)** Comparison of shared Matrisome peptides (PSMs) identified in each fraction. A paired two-tailed Student t test was used to compare means between groups. The number of peptide identifications made in each sub-category was significantly higher in the Azo fraction than the Decell fraction. In the Decell fraction, 320 collagen peptides were identified and in the Azo fraction, 1,101 collagen peptides were identified (*p* = 2.8E-3). A total of 1,417 glycoprotein peptides were identified in the Decell fraction and 3,071 peptides were identified in the Azo fraction (*p* = 0.010). Finally, 220 proteoglycan peptides were identified in the Decell fraction and 533 peptides were identified in the Azo fraction (*p* = 0.0035). Levels of statistical significance are notated with an asterisk (*): \**p* ≤ 0.05, \*\**p* < 0.01, and \*\*\**p* < 0.001; no statistical significance *(ns)* if *p* > 0.05. **C)** Comparison of Matrisome peptides unique to each fraction. A total of 1,358 unique collagen peptides were identified between the Decell and Azo fractions. 983 collagen peptides were uniquely identified in the Azo fraction and 18 peptides in the Decell fraction. 357 peptides were identified in both fractions. A total of 4,021 proteoglycan matrisome peptide identifications were made, with 194 peptides unique in the Decell fraction and 2,406 unique to the Azo fraction, and 1,421 peptides shared in both fractions. Finally, a total of 656 unique proteoglycan peptide identifications were made between the two fractions. 8 unique peptides were identified in the Azo fraction and 408 were unique to the Azo fraction, with 240 peptides in common. **D)** Comparison of intensities of representative core matrisome peptides shared between the Decell and Azo fractions. The most abundant peptide in the Decell was selected for COL6A3 (Collagen alpha-3(VI) chain), FN1 (Fibronectin, a key glycoprotein), and OGN (Mimecan, a proteoglycan). A paired two-tailed Student t test was used to determine compare means between groups. Levels of statistical significance are notated with an asterisk (*): \**p* ≤ 0.05, \*\**p* < 0.01, and \*\*\**p* < 0.001; no statistical significance *(ns)* if *p* > 0.05. **E)** The insoluble ECM (iECM) is recovered in the Azo fraction. 8 Fibril-forming collagen species were consistently recovered in the Azo fraction, with protein intensities ranging from 2-8% RSD. **F)** Protein intensities of alternatively-spliced Elastin isoforms. Isoform 1 was identified in both fractions, and isoforms 5 and 11 were recovered in the Azo fraction. **G)** Average percent sequence coverage for iECM proteins shown in panels E and F. Average (±SE) sequence coverage for fibrillary collagens was 26.7 ±0.8% for the 8 species in panel E, and the average sequence coverage for the 3 alternatively spliced isoforms of Elastin was 15.57 ±0.1%.

On the peptide level, significantly higher numbers of core matrisome peptides were identified in the Azo fraction compared to the Decell fraction (**Figure 3B**). We identified 1,101 unique collagen peptides in the Azo fraction compared to 320 peptides in the Decell (*p* = 2.8E-3, paired two-tailed t test). A total of 3,071 peptides related to glycoproteins were identified in the Azo fraction compared to 1,417 peptides in the Decell (*p* = 0.010). Finally, 533 proteoglycan peptides were identified in the Azo fraction compared to 220 peptides in the Decell fraction (*p* = 0.0035). In total, we identified an impressive 1,358 collagen peptides, 4,021 glycoprotein peptides, and 656 proteoglycan peptides between the two extracts (**Figure 3C**).

The majority of core matrisome peptides were identified solely in the Azo fraction and very few core matrisome peptides were exclusively identified in the Decell fraction (**Figure 3C**). Of the total peptides identified, 72.4% of collagen peptides, 59.8% of glycoprotein peptides, and 62.2% of proteoglycan peptides were identified exclusively in Azo, while only 1.3%, 4.8%, and 1.2% were identified solely in Decell (**Table S2**). Among the peptides identified in both extracts, peptide abundance was significantly higher in the Azo extract. The highest intensity peptides corresponding to a prominent core matrisome proteins in the Decell fraction were selected and compared to the peptide detected in the Azo fraction. For each comparison, the peptide intensity was significantly higher in the Azo fraction compared to the Decell fraction (**Figure 3D**). For example, 108 peptides corresponding to Collagen alpha-3(VI) chain (COL6A3) were identified in the Decell fraction, and 369 peptides were identified in the Azo fraction. The most abundant peptide in the Decell, LSDAGITPLFLTR2, was significantly lower in abundance compared to the same peptide found in the Azo extract (*p* = 0.014, two-tailed paired t test). Similar results were found for peptides corresponding to prominent glycoprotein Fibronectin (gene: FN1, *p* = 0.0052) and proteoglycan Mimecan (gene: OGN, *p* = 0.036). Thus, Azo significantly enriches core matrisome proteins.

In the Azo fraction, we achieved excellent recovery of the insoluble matrisome, defined as the dense, fibrous, cross-linked proteins essential for maintaining lung architecture and elasticity. Importantly, we detected 1,072 unique peptides corresponding to all 32 unique collagen isoforms and all 28 collagen species known in humans (summarized in **Table S4**). Fibrillar collagens I, II, III, V, and XI^13^ were recovered, with protein intensities spanning 1-4% RSD across extraction replicates (**Figure 3E**). We reproducibly recovered three alternatively-spliced isoforms of elastin, part of the insoluble ECM that is responsible for the expansion capability of the lung (**Figure 3F**).^37, 38^ Isoform 1 was identified in both fractions, and isoforms 5 and 11 were recovered exclusively in the Azo fraction. Average sequence coverage for the 8 fibrillar collagens detected was 26.7 ±0.8% and 15.6 ±0.1% for three elastin isoforms (**Figure 3G**).

### Matrisome-Associated Proteins are Found in Both Extracts

Matrisome Associated proteins, including ECM-affiliated proteins, ECM regulators, and secreted factors were identified in both fractions. In total, 249 matrisome associated protein groups were identified between the Azo and Decell extracts, with approximately 51% of identifications shared between fractions (**Figure 4A**). No significant difference in peptide count was found between the Decell and Azo extracts for ECM-affiliated proteins and ECM regulators, indicating no specific enrichment in a sample fraction (**Figure 4B**). In the Decell fraction, 244 peptides matching to secreted factors and 180 peptides in the Azo fraction were identified (*p* = 0.017). Peptide overlap between the Decell and Azo extracts was 50% for ECM-affiliated peptides, 38% for secreted factors, and 20% for ECM regulators, with larger proportions of matrisome associated peptides exclusively identified in each fraction (**Figure 4C**).

**Figure 4.**
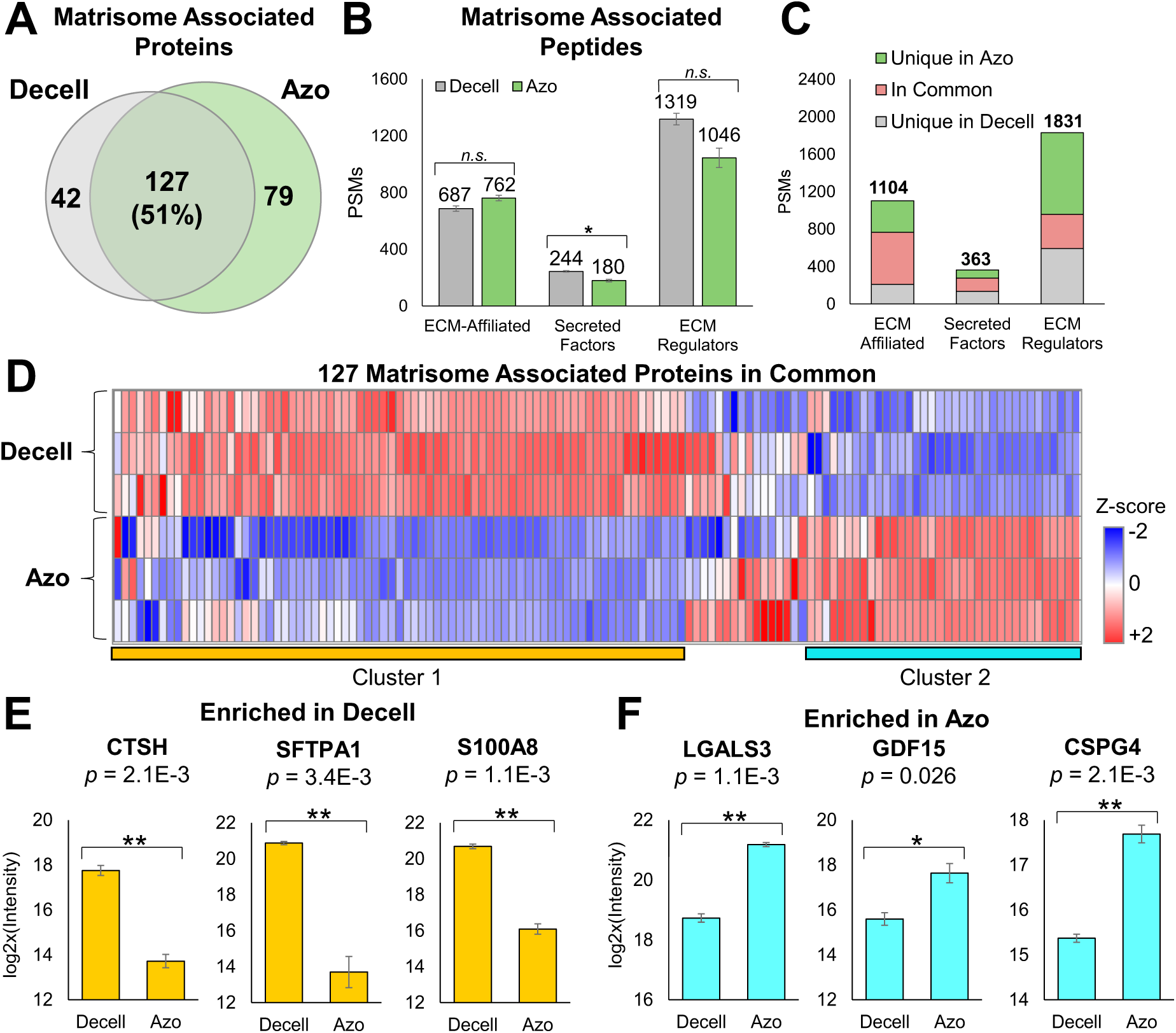
Matrisome-associated proteins are identified in both fractions. **A)** Comparison of Matrisome Associated protein group identifications in the Decell and Azo Fractions. 43 protein groups were unique to the Decell fraction, and 80 proteins were unique to the Azo fraction. 126 proteins (51% of total IDs) were identified in both fractions. **B)** Comparison of Matrisome peptides (PSMs) identified in each fraction. In the Decell fraction, 687 ECM-Affiliated proteins were identified and in the Azo, fraction, 762 peptides were identified (*p* = 0.11). In the Decell fraction, 244 peptides matching Secreted Factors and 180 peptides in the Azo fraction were identified (*p* = 0.017). 1,319 ECM Regulator peptides were identified in the Decell 1,046 peptides were identified in the Azo fraction (*p* = 0.073). A paired two-tailed Student t test was used to compare means between groups. Levels of statistical significance are notated with an asterisk (*): \**p* ≤ 0.05, \*\**p* < 0.01, and \*\*\**p* < 0.001; no statistical significance *(ns)* if *p* > 0.05. **C)** Comparison of Matrisome-Associated peptides unique to each fraction. A total of 1,104 unique ECM-Affiliated peptides were identified between the Decell and Azo fractions combined. 210 peptides (19% of total) were identified exclusively in the Decell fraction, and 337 (31% of total) ECM-Affiliated peptides were uniquely identified in the Azo fraction. 557 peptides were identified in both fractions. A total of 363 unique secreted factors peptides were identified, and 136 peptides (37% of total) were unique to the Decell fraction and 88 (24%) were unique to the Azo fraction. 139 peptides were shared between fractions. Finally, a total of 1,831 unique ECM Regulator peptides were identified between two fractions. 593 (32% of total) unique peptides were identified in the Decell fraction and 874 (48%) peptides were unique to the Azo fraction, and 364 peptides were identified in both fractions. **D)** Differential enrichment of the 126 shared Matrisome-Associated protein identifications between Decell and Azo fractions. Z-Score normalization was performed on log2x-transfrmed protein intensities prior to unsupervised hierarchical clustering. Clusters 1 and 2 highlight the most different protein intensities between the two fractions. Cluster 1 represents proteins most abundant in the Decell fraction, and Cluster 2 represents proteins most abundant in the Azo fraction. **E)** Comparison of protein intensities of representative Matrisome-associated proteins enriched in the Decell fraction included in Cluster 1 in panel **D**, including Pro-cathepsin H (gene: CTSH), Pulmonary surfactant-associated protein A1 (gene: SFTPA1), and Protein S100-A8 (gene: S100A8). Gene names are reported above each plot and significance values were determined by two-sample t test; Benjamini-Hotchberg corrected *p* values are reported. Levels of statistical significance are notated with an asterisk (*): \**p* ≤ 0.05, \*\**p* < 0.01, and \*\*\**p* < 0.001; no statistical significance *(ns)* if *p* > 0.05. **F)** Comparison of protein intensities of representative Matrisome-associated proteins enriched in the Azo fraction, highlighted as Cluster 2 in panel **D**, including Galectin-3 (gene: LGALS3), Growth/differentiation factor 15 (gene: GDF15), and Chondroitin sulfate proteoglycan 4 (gene: CSPG4).

Of 126 shared matrisome associated proteins between the two fractions, 89 proteins were enriched on the protein level (two-sample t test, *p*<0.05, Benjamini-Hotchberg correction). 57 proteins were significantly enriched in the Decell fraction and 32 proteins were significantly enriched in the Azo fraction (**Figure 4D**). Proteins shown to be important to ECM regulation including Pro-cathepsin H (gene: CTSH), Pulmonary surfactant-associated protein A1 (gene: SFTPA1), and Protein S100-A8 (gene: S100A8) were enriched in the Decell fraction (**Figure 4E**). Additionally, five cathepsin proteins were enriched in this fraction, including Cathepsin D (CTSD), Pro-cathepsin H (CTSH), Procathepsin L (CTSL), Cathepsin S (CTSS), and Cathepsin Z (CTSZ). Pulmonary surfactant-associated protein A1 (SFTPA1), a secreted protein reducing tension on the alveolar surface,^39^ was also enriched in the Decell fraction. Additionally, several S100 proteins were enriched in the Decell fraction, which are secreted factors whose dysregulation has been linked to lung disease.^40^

Matrisome associated proteins enriched in the Azo fraction included Galectin-3 (gene: LGALS3), growth/differentiation factor 15 (gene: GDF15), and Chondroitin sulfate proteoglycan 4 (gene: CSPG4) (**Figure 4F**). Included in the proteins enriched in this fraction were four Galectin species (genes LGALS1, LGALS3, LGALS8, and LGALS9), which have been found to regulate transforming growth factor beta-1 pathways in idiopathic pulmonary fibrosis (IPF).^41^ Growth factors associated with ECM synthesis and dysregulation in IPF^42^ including growth/differentiation factor 15 (GDF15) and transforming growth factor beta-1 proprotein (TGFB1) were enriched in the Azo fraction. Additionally, proteoglycan-binding proteins such as Chondroitin sulfate proteoglycan 4 (CSPG4)^43^ were enriched in this fraction. Protein classes such as procollagen proteins involved in lysine hydroxylation, including Prolyl 3-hydroxylase 1 (P3H1), Multifunctional procollagen lysine (PLOD3), and Isoform 2 of Prolyl 4-hydroxylase subunit alpha-1 (P4HA1).

### Overview of Pulmonary Extracellular Matrix Coverage

Overall, we have identified 392 unique matrisome genes and 418 alternatively-spliced matrisome isoforms. Key components of the basement membrane and interstitial ECM in human lungs are identified in this dataset, including basement membrane proteins such as integrins, laminins, nidogen, perlecan, and collagens IV, as well as the interstitial matrix including elastin, fibrillar collagen species I, II, III, V, and XI, proteoglycans, and fibronectin (**Figure 5A**). The matrisome coverage this method achieves allows for a comprehensive view of protein network and pathways prominent in the human lung ECM. For example, 133 proteins are associated with extracellular matrix organization (Reactome Pathway HSA-1474244, FDR-corrected *p* = 1.47E-119), and 51 IDs are associated with collagen formation (Reactome Pathway HSA-1474290, FDR-corrected *p* = 3.25E-47) (**Figure 5B**). Protein-protein interactions within the dataset include elastic fiber formation (**Figure 5C**), and laminin interactions (**Figure 5D**). We envision this discovery-based ECM proteomics method can be used to identify potential therapeutic targets and advance understanding of ECM-mediated lung pathogenesis.

**Figure 5.**
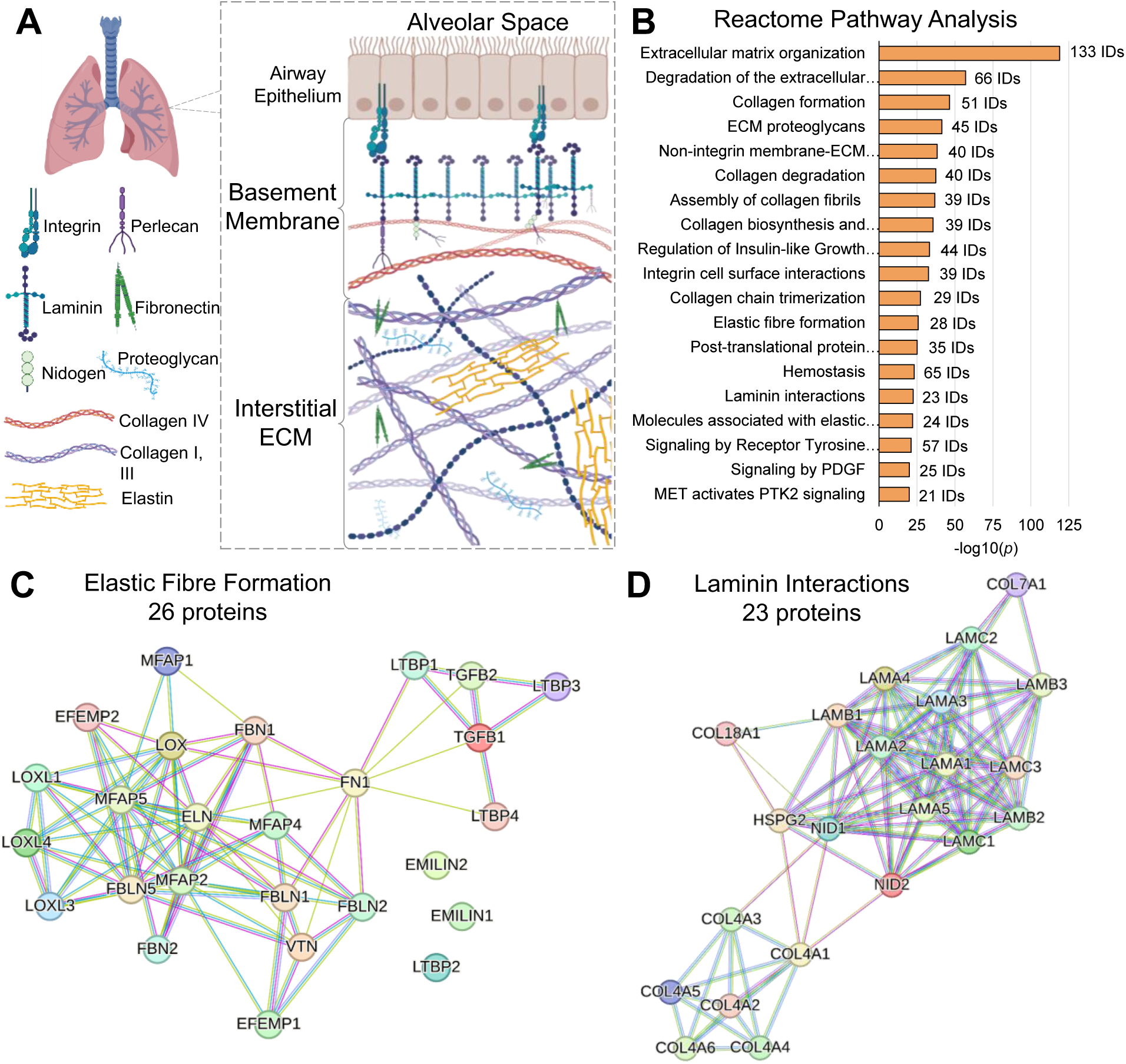
Overview of Pulmonary Extracellular Matrix Coverage. **A)** Major components of the basement membrane and interstitial ECM were detected in our data, including basement membrane proteins, integrins, laminins, nidogen, Perlecan, collagens IV. Proteins comprising the interstitial matrix such as elastin, fibrillar collagen species I, II, III, V, and XI, proteoglycans, and fibronectin were identified. **B)** Top 20 most significantly enriched Reactome Pathways resulting from STRING Protein-Protein Interaction Analysis. Pathways are sorted by –log10(FDR-adjusted *p* value), and gene count is provided next to each bar. **C)** String interactome map of Reactome Pathway HSA-1566948, Elastic Fibre Formation (FDR-corrected *p* = 1.70E-26), showing protein-protein interaction network built with identified matrisome proteins. STRING analysis was performed with high (<0.7) confidence, and connections between nodes are evidence based (textmining, databases, and experimentally-determined interaction**s**). **D)** STRING interactome map showing Reactome Pathway HSA-3000157, Laminin Interactors (FDR-corrected *p* = 3.34E-23), with identified proteins from our data. STRING analysis was performed with high (<0.7) confidence, and connections between nodes are evidence based (textmining, databases, and experimentally-determined interaction**s**). Overrepresentation was determined based on background gene list.

## Conclusion

Overall, this methodology enables reproducible and rapid access to the pulmonary ECM from human lungs. The single-step solubilization of the core matrisome using Azo enables a facile means of understanding of the highly complex 3D architecture of lung tissue. We have demonstrated the successful enrichment of core matrisome proteins including collagens, proteoglycans, and glycoproteins using Azo. This discovery-based lung ECM proteomics strategy can be used to generate hypotheses and identify potential therapeutic targets, and advance understanding of extracellular-mediated lung pathogenesis.

## Acknowledgements

We are grateful to the donors and their families for their generous donation of tissue. We thank James Anderson and Carrie Sparks at the University of Wisconsin Organ and Tissue Donation for coordination of donor lung collection. We thank those who assisted with pulmonary tissue collection and dissection. AbbVie funded the study and participated in study design, research, data collection, analysis and interpretation of data, writing, reviewing, and approving the publication. We acknowledge PNNL and the Protein-Coverage-Summarizer GitHub repository for use of the Sequence Coverage Summarizer tool.

## Declarations of interest

Funding for this work was provided by AbbVie, Inc.

V.M.T., S.L., Y.H., and Y.T. are employees of AbbVie, Inc. AbbVie participated in the design and conduct of this research. AbbVie participated in the interpretation of data, review, and approval of the publication.

AbbVie takes no position on the tools and technologies shown. All statements regarding tools and technologies are the personal opinion of the authors.

## Data Availability

Raw data are available in the MassIVE repository with identifier MSV000092235.

## References

(1) Mouw, J. K.; Ou, G.; Weaver, V. M. Extracellular matrix assembly: a multiscale deconstruction. Nature Reviews Molecular Cell Biology 2014, 15 (12), 771–785. DOI: 10.1038/nrm3902. Zhou, Y.; Horowitz, J. C.; Naba, A.; Ambalavanan, N.; Atabai, K.; Balestrini, J.; Bitterman, P. B.; Corley, R. A.; Ding, B.-S.; Engler, A. J.;, et al. Extracellular matrix in lung development, homeostasis and disease. Matrix Biology 2018, 73, 77–104. DOI: https://doi.org/10.1016/j.matbio.2018.03.005.

(2) Daley, W. P.; Peters, S. B.; Larsen, M. Extracellular matrix dynamics in development and regenerative medicine. Journal of Cell Science 2008, 121 (3), 255–264. DOI: 10.1242/jcs.006064 (acccessed 6/22/2023).

(3) Hannu, J.; Annele, S.; Markku, K.; Thomas, N. W.; Risto, P. Extracellular Matrix Molecules: Potential Targets in Pharmacotherapy. Pharmacological Reviews 2009, 61 (2), 198. DOI: 10.1124/pr.109.001289. Gerald, B.; Bettina, O.; Michael, G.; Eric, S. W.; Herbert, B. S.; Oliver, E. The instructive extracellular matrix of the lung: basic composition and alterations in chronic lung disease. European Respiratory Journal 2017, 50 (1), 1601805. DOI: 10.1183/13993003.01805-2016.

(4) Burgess, J. K.; Mauad, T.; Tjin, G.; Karlsson, J. C.; Westergren-Thorsson, G. The extracellular matrix – the under-recognized element in lung disease? The Journal of Pathology 2016, 240 (4), 397–409, https://doi.org/10.1002/path.4808. DOI: https://doi.org/10.1002/path.4808 (acccessed 2023/06/18).

(5) Fernandez, I. E.; Eickelberg, O. New cellular and molecular mechanisms of lung injury and fibrosis in idiopathic pulmonary fibrosis. The Lancet 2012, 380 (9842), 680–688. DOI: 10.1016/S0140-6736(12)61144-1 (acccessed 2023/06/21). Watson, W. H.; Ritzenthaler, J. D.; Roman, J. Lung extracellular matrix and redox regulation. Redox Biology 2016, 8, 305–315. DOI: https://doi.org/10.1016/j.redox.2016.02.005.

(6) Cromar, G. L.; Xiong, X.; Chautard, E.; Ricard-Blum, S.; Parkinson, J. Toward a systems level view of the ECM and related proteins: A framework for the systematic definition and analysis of biological systems. Proteins: Structure, Function, and Bioinformatics 2012, 80 (6), 1522–1544, https://doi.org/10.1002/prot.24036. DOI: https://doi.org/10.1002/prot.24036 (acccessed 2023/06/21).

(7) Naba, A. Ten Years of Extracellular Matrix Proteomics: Accomplishments, Challenges, and Future Perspectives. Molecular & Cellular Proteomics 2023, 22 (4). DOI: 10.1016/j.mcpro.2023.100528 (acccessed 2023/06/18).

(8) Naba, A.; Clauser, K. R.; Hoersch, S.; Liu, H.; Carr, S. A.; Hynes, R. O. The Matrisome: In Silico Definition and In Vivo Characterization by Proteomics of Normal and Tumor Extracellular Matrices. Molecular & Cellular Proteomics 2012, 11 (4). DOI: 10.1074/mcp.M111.014647 (acccessed 2023/06/18).

(9) Naba, A.; Hoersch, S.; Hynes, R. O. Towards definition of an ECM parts list: An advance on GO categories. Matrix Biology 2012, 31 (7), 371–372. DOI: https://doi.org/10.1016/j.matbio.2012.11.008.

(10) Shao, X.; Gomez, C. D.; Kapoor, N.; Considine, J. M.; Grams, C.; Gao, Y.; Naba, A. MatrisomeDB 2.0: 2023 updates to the ECM-protein knowledge database. Nucleic Acids Research 2023, 51 (D1), D1519–D1530. DOI: 10.1093/nar/gkac1009 (acccessed 3/22/2023).

(11) Krasny, L.; Bland, P.; Kogata, N.; Wai, P.; Howard, B. A.; Natrajan, R. C.; Huang, P. H. SWATH mass spectrometry as a tool for quantitative profiling of the matrisome. Journal of Proteomics 2018, 189, 11–22. DOI: https://doi.org/10.1016/j.jprot.2018.02.026.

(12) Barrett, A. S.; Wither, M. J.; Hill, R. C.; Dzieciatkowska, M.; D’Alessandro, A.; Reisz, J. A.; Hansen, K. C. Hydroxylamine Chemical Digestion for Insoluble Extracellular Matrix Characterization. Journal of Proteome Research 2017, 16 (11), 4177–4184. DOI: 10.1021/acs.jproteome.7b00527.

(13) Bella, J.; Hulmes, D. J. S. Fibrillar Collagens. In Fibrous Proteins: Structures and Mechanisms, Parry, D. A. D., Squire, J. M. Eds.; Springer International Publishing, 2017; pp 457–490.

(14) Naba, A.; Clauser, K. R.; Ding, H.; Whittaker, C. A.; Carr, S. A.; Hynes, R. O. The extracellular matrix: Tools and insights for the “omics” era. Matrix Biology 2016, 49, 10–24. DOI: https://doi.org/10.1016/j.matbio.2015.06.003.

(15) Naba, A. Molecular & Cellular Proteomics 2023, *22* (4). DOI: 10.1016/j.mcpro.2023.100528 (acccessed 2023/06/18).

(16) Naba, A.; Pearce, O. M. T.; Del Rosario, A.; Ma, D.; Ding, H.; Rajeeve, V.; Cutillas, P. R.; Balkwill, F. R.; Hynes, R. O. Characterization of the Extracellular Matrix of Normal and Diseased Tissues Using Proteomics. Journal of Proteome Research 2017, 16 (8), 3083–3091. DOI: 10.1021/acs.jproteome.7b00191. Hill, R. C.; Calle, E. A.; Dzieciatkowska, M.; Niklason, L. E.; Hansen, K. C. Quantification of Extracellular Matrix Proteins from a Rat Lung Scaffold to Provide a Molecular Readout for Tissue Engineering*[S]. Molecular & Cellular Proteomics 2015, 14 (4), 961–973. DOI: https://doi.org/10.1074/mcp.M114.045260.

(17) Schiller, H. B.; Fernandez, I. E.; Burgstaller, G.; Schaab, C.; Scheltema, R. A.; Schwarzmayr, T.; Strom, T. M.; Eickelberg, O.; Mann, M. Time- and compartment-resolved proteome profiling of the extracellular niche in lung injury and repair. Molecular Systems Biology 2015, 11 (7), 819, https://doi.org/10.15252/msb.20156123. DOI: https://doi.org/10.15252/msb.20156123 (acccessed 2023/06/21). McCabe, M. C.; Schmitt, L. R.; Hill, R. C.; Dzieciatkowska, M.; Maslanka, M.; Daamen, W. F.; van Kuppevelt, T. H.; Hof, D. J.; Hansen, K. C. Evaluation and Refinement of Sample Preparation Methods for Extracellular Matrix Proteome Coverage. Molecular & Cellular Proteomics 2021, 20. DOI: 10.1016/j.mcpro.2021.100079 (acccessed 2022/04/21).

(18) McCabe, M. C.; Saviola, A. J.; Hansen, K. C. Mass Spectrometry-Based Atlas of Extracellular Matrix Proteins across 25 Mouse Organs. Journal of Proteome Research 2023, 22 (3), 790–801. DOI: 10.1021/acs.jproteome.2c00526.

(19) Au - Naba, A.; Au - Clauser, K. R.; Au - Hynes, R. O. Enrichment of Extracellular Matrix Proteins from Tissues and Digestion into Peptides for Mass Spectrometry Analysis. JoVE 2015, (101), e53057. DOI: doi:10.3791/53057.

(20) Brown, K. A.; Chen, B.; Guardado-Alvarez, T. M.; Lin, Z.; Hwang, L.; Ayaz-Guner, S.; Jin, S.; Ge, Y. A photocleavable surfactant for top-down proteomics. Nature Methods 2019, 16 (5), 417–420. DOI: 10.1038/s41592-019-0391-1.

(21) Meier, F.; Brunner, A.-D.; Frank, M.; Ha, A.; Bludau, I.; Voytik, E.; Kaspar-Schoenefeld, S.; Lubeck, M.; Raether, O.; Bache, N.;, et al. diaPASEF: parallel accumulation–serial fragmentation combined with data-independent acquisition. Nature Methods 2020, 17 (12), 1229–1236. DOI: 10.1038/s41592-020-00998-0.

(22) Brown, K. A.; Tucholski, T.; Eken, C.; Knott, S.; Zhu, Y.; Jin, S.; Ge, Y. High-Throughput Proteomics Enabled by a Photocleavable Surfactant. Angewandte Chemie International Edition 2020, 59 (22), 8406–8410, https://doi.org/10.1002/anie.201915374. DOI: https://doi.org/10.1002/anie.201915374 (acccessed 2021/01/18).

(23) Aballo, T. J.; Roberts, D. S.; Melby, J. A.; Buck, K. M.; Brown, K. A.; Ge, Y. Ultrafast and Reproducible Proteomics from Small Amounts of Heart Tissue Enabled by Azo and timsTOF Pro. Journal of Proteome Research 2021, 20 (8), 4203–4211. DOI: 10.1021/acs.jproteome.1c00446.

(24) Knott, S. J.; Brown, K. A.; Josyer, H.; Carr, A.; Inman, D.; Jin, S.; Friedl, A.; Ponik, S. M.; Ge, Y. Photocleavable Surfactant-Enabled Extracellular Matrix Proteomics. Analytical Chemistry 2020, 92 (24), 15693–15698. DOI: 10.1021/acs.analchem.0c03104.

(25) Meier, F.; Brunner, A.-D.; Koch, S.; Koch, H.; Lubeck, M.; Krause, M.; Goedecke, N.; Decker, J.; Kosinski, T.; Park, M. A.;, et al. Online Parallel Accumulation–Serial Fragmentation (PASEF) with a Novel Trapped Ion Mobility Mass Spectrometer. Molecular & Cellular Proteomics 2018, 17 (12), 2534–2545. DOI: 10.1074/mcp.TIR118.000900 (acccessed 2021/03/05).

(26) Demichev, V.; Szyrwiel, L.; Yu, F.; Teo, G. C.; Rosenberger, G.; Niewienda, A.; Ludwig, D.; Decker, J.; Kaspar-Schoenefeld, S.; Lilley, K. S.; et al. dia-PASEF data analysis using FragPipe and DIA-NN for deep proteomics of low sample amounts. Nature Communications 2022, 13 (1), 3944. DOI: 10.1038/s41467-022-31492-0.

(27) Demichev, V.; Messner, C. B.; Vernardis, S. I.; Lilley, K. S.; Ralser, M. DIA-NN: neural networks and interference correction enable deep proteome coverage in high throughput. Nature Methods 2020, 17 (1), 41–44. DOI: 10.1038/s41592-019-0638-x.

(28) Hulsen, T.; de Vlieg, J.; Alkema, W. BioVenn – a web application for the comparison and visualization of biological lists using area-proportional Venn diagrams. BMC Genomics 2008, 9 (1), 488. DOI: 10.1186/1471-2164-9-488.

(29) Wiśniewski, J. R. Chapter Four - Label-Free and Standard-Free Absolute Quantitative Proteomics Using the “Total Protein” and “Proteomic Ruler” Approaches. In Methods in Enzymology, Shukla, A. K. Ed.; Vol. 585; Academic Press, 2017; pp 49–60. Wiśniewski, J. R.; Ostasiewicz, P.; Duś, K.; Zielińska, D. F.; Gnad, F.; Mann, M. Extensive quantitative remodeling of the proteome between normal colon tissue and adenocarcinoma. Molecular Systems Biology 2012, 8 (1), 611, https://doi.org/10.1038/msb.2012.44. DOI: https://doi.org/10.1038/msb.2012.44 (acccessed 2023/06/21).

(30) Huang, D. W.; Sherman, B. T.; Lempicki, R. A. Systematic and integrative analysis of large gene lists using DAVID bioinformatics resources. Nature Protocols 2009, 4 (1), 44–57. DOI: 10.1038/nprot.2008.211.

(31) Thomas, P. D.; Ebert, D.; Muruganujan, A.; Mushayahama, T.; Albou, L.-P.; Mi, H. PANTHER: Making genome-scale phylogenetics accessible to all. Protein Science 2022, 31 (1), 8–22, https://doi.org/10.1002/pro.4218. DOI: https://doi.org/10.1002/pro.4218 (acccessed 2023/04/18).

(32) Szklarczyk, D.; Kirsch, R.; Koutrouli, M.; Nastou, K.; Mehryary, F.; Hachilif, R.; Gable, A. L.; Fang, T.; Doncheva, Nadezhda T.; Pyysalo, S.;, et al. The STRING database in 2023: protein– protein association networks and functional enrichment analyses for any sequenced genome of interest. Nucleic Acids Research 2023, 51 (D1), D638–D646. DOI: 10.1093/nar/gkac1000 (acccessed 6/21/2023).

(33) Yeung, Y.-G.; Nieves, E.; Angeletti, R. H.; Stanley, E. R. Removal of detergents from protein digests for mass spectrometry analysis. Analytical Biochemistry 2008, 382 (2), 135–137. DOI: https://doi.org/10.1016/j.ab.2008.07.034.

(34) Starcher, B. C. Lung Elastin and Matrix. CHEST 2000, 117 (5), 229S–234S. DOI: 10.1378/chest.117.5_suppl_1.229S-a (acccessed 2023/06/23).

(35) Naba, A.; Clauser, K. R.; Hoersch, S.; Liu, H.; Carr, S. A.; Hynes, R. O. The Matrisome: *In Silico* Definition and *In Vivo* Characterization by Proteomics of Normal and Tumor Extracellular Matrices *. Molecular & Cellular Proteomics 2012, 11 (4). DOI: 10.1074/mcp.M111.014647 (acccessed 2023/06/18).

(36) Lansky, Z.; Mutsafi, Y.; Houben, L.; Ilani, T.; Armony, G.; Wolf, S. G.; Fass, D. 3D mapping of native extracellular matrix reveals cellular responses to the microenvironment. Journal of Structural Biology: X 2019, 1, 100002. DOI: https://doi.org/10.1016/j.yjsbx.2018.100002.

(37) Vindin, H. J.; Oliver, B. G. G.; Weiss, A. S. Elastin in healthy and diseased lung. Current Opinion in Biotechnology 2022, 74, 15–20. DOI: https://doi.org/10.1016/j.copbio.2021.10.025.

(38) Wagenseil, J. E.; Mecham, R. P. New insights into elastic fiber assembly. Birth Defects Research Part C: Embryo Today: Reviews 2007, 81 (4), 229–240, https://doi.org/10.1002/bdrc.20111. DOI: https://doi.org/10.1002/bdrc.20111 (acccessed 2023/07/18).

(39) Han, S.; Mallampalli, R. K. The Role of Surfactant in Lung Disease and Host Defense against Pulmonary Infections. Annals of the American Thoracic Society 2015, 12 (5), 765–774. DOI: 10.1513/AnnalsATS.201411-507FR (acccessed 2023/06/21).

(40) Sattar, Z.; Lora, A.; Jundi, B.; Railwah, C.; Geraghty, P. The S100 Protein Family as Players and Therapeutic Targets in Pulmonary Diseases. Pulmonary Medicine 2021, 2021, 5488591. DOI: 10.1155/2021/5488591.

(41) MacKinnon, A. C.; Gibbons, M. A.; Farnworth, S. L.; Leffler, H.; Nilsson, U. J.; Delaine, T.; Simpson, A. J.; Forbes, S. J.; Hirani, N.; Gauldie, J.;, et al. Regulation of Transforming Growth Factor-β1–driven Lung Fibrosis by Galectin-3. American Journal of Respiratory and Critical Care Medicine 2012, 185 (5), 537–546. DOI: 10.1164/rccm.201106-0965OC (acccessed 2023/06/21). Zick, Y.; Eisenstein, M.; Goren, R. A.; Hadari, Y. R.; Levy, Y.; Ronen, D. Role of galectin-8 as a modulator of cell adhesion and cell growth. Glycoconjugate Journal 2002, 19 (7), 517–526. DOI: 10.1023/B:GLYC.0000014081.55445.af.

(42) Radwanska, A.; Cottage, C. T.; Piras, A.; Overed-Sayer, C.; Sihlbom, C.; Budida, R.; Wrench, C.; Connor, J.; Monkley, S.; Hazon, P.;, et al. Increased expression and accumulation of GDF15 in IPF extracellular matrix contribute to fibrosis. JCI Insight 2022, 7 (16). DOI: 10.1172/jci.insight.153058. Saito, A.; Horie, M.; Nagase, T. TGF-β Signaling in Lung Health and Disease. In International Journal of Molecular Sciences, 2018; Vol. 19.

(43) Shannon, J. M.; McCormick-Shannon, K.; Burhans, M. S.; Shangguan, X.; Srivastava, K.; Hyatt, B. A. Chondroitin sulfate proteoglycans are required for lung growth and morphogenesis in vitro. American Journal of Physiology-Lung Cellular and Molecular Physiology 2003, 285 (6), L1323–L1336. DOI: 10.1152/ajplung.00226.2003 (acccessed 2023/06/21).

