## Supplemental Information for "High-throughput Extracellular Matrix Proteomics of Human Lungs Enabled by Photocleavable Surfactant and diaPASEF"

**Table of Contents**

**Supplemental Tables**

- Table S1. Donor clinical characteristics of n=3 lung provided by the UW Hospital Organ Procurement Organization.
- Table S2. Core Matrisome Peptides Comparison Between Decell and Azo Extracts.
- Table S3. Core Matrisome Proteins Identified Exclusively in Azo Fraction.
- Table S4. Summary of Collagen Species detected in Azo Fraction

**Supplemental Figures**

- Figure S1. DPBS Wash Post-Cryopulverization Protein Increases Total Protein Signal by Depleting Plasma Proteins.
- Figure S2. Comparison of HEPES and Triton X-100 based Decellularization of Lung Tissue
- Figure S3. Strong Reproducibility between Extraction Replicates Observed for all Identified Proteins.
- Figure S4. Global Overview of Decell and Azo Fraction Content.
- Figure S5. Overview of Matrisome Recovery in the Decell Fraction.
- Figure S6. Comparison of core matrisome identifications by Azo the predicted and experimentally observed matrisome.

**Table S1.** **Donor clinical characteristics of n=3 donor lungs provided by the UW Hospital Organ Procurement Organization.** All donors had no history of pulmonary disease and were not used for transplant due to antigen incompatibility, transplant candidate availability, or co-morbidity. Average and Standard Error (SE) is reported for age. Number of donors (n) is indicated in parentheses after each reported percentage.

| **Clinical Characteristic** | **Percentage of sample cohort** |
| --- | --- |
| Age | 38 ± 1.7 years |
|  | min: 35 years |
|  | max: 41 years |
| Male | 67% (n = 2) |
| Female | 33% (n = 1) |
| Asthma | 33% (n = 1) |
| Substance Abuse | 67% (n = 2) |
| Smoking | 67% (n = 2) |

**Table S2. Core Matrisome Peptides Comparison Between Decell and Azo Extracts.** Total number of core matrisome peptides, PSMs, that are exclusively identified in each fraction and shared between two fractions (Decell and Azo) and the relative percentage of the total amount of PSMs identified for each category.

|  | **Unique in Decell** | | **Unique in Azo** | | **Shared** | |
| --- | --- | --- | --- | --- | --- | --- |
| **Core Matrisome** | **PSMs** | **% of Total** | **PSMs** | **%** | **PSMs** | **%** |
| Collagens | 18 | 1.3% | 983 | 72% | 357 | 26% |
| Glycoproteins | 194 | 4.8% | 2,406 | 60% | 1,421 | 35% |
| Proteoglycans | 8 | 1.2% | 408 | 62% | 240 | 37% |

**Table S3. Core Matrisome Proteins Identified Exclusively in Azo Fraction.** List of core matrisome proteins that were only identified in the Azo Fraction, including UniProt Accession number (UniProt ID), Gene, and Protein Name, and approximate concentration in solution estimated by Total Protein Analysis.^1^ Average and Standard Error (SE) between n=3 extraction replicates is reported.

| **Protein Name** | **UniProt ID** | **Gene** | **Avg (pmol/mg)** | **SE** |
| --- | --- | --- | --- | --- |
| Neurocan core protein | O14594;Q96GW7;Q96GW7-2 | NCAN | 0.20 | 0.11 |
| BMP-binding endothelial regulator protein | Q8N8U9 | BMPER | 0.12 | 0.02 |
| Collagen alpha-1(X) chain | Q03692 | COL10A1 | 0.30 | 0.02 |
| Collagen alpha-1(XI) chain | P12107;P12107-2;P12107-3;P12107-4 | COL11A1 | 0.08 | 0.02 |
| Collagen alpha-1(XVI) chain | Q07092;Q07092-2 | COL16A1 | 0.19 | 0.02 |
| Collagen alpha-1(XXI) chain | Q96P44;Q96P44-2;Q96P44-3 | COL21A1 | 4.30 | 0.58 |
| Collagen alpha-1(XXVIII) chain | Q2UY09 | COL28A1 | 0.36 | 0.03 |
| Collagen alpha-1(II) chain | P02458;P02458-1 | COL2A1 | 0.04 | 0.01 |
| Collagen alpha-3(IV) chain | Q01955 | COL4A3 | 2.30 | 0.99 |
| Collagen alpha-4(IV) chain | P53420 | COL4A4 | 1.45 | 0.14 |
| Collagen alpha-5(IV) chain | P29400;P29400-2 | COL4A5 | 0.27 | 0.04 |
| Collagen alpha-6(IV) chain | Q14031;Q14031-2 | COL4A6 | 0.02 | 0.02 |
| Collagen alpha-2(V) chain | P05997 | COL5A2 | 0.60 | 0.04 |
| Collagen alpha-3(V) chain | P25940 | COL5A3 | 1.23 | 0.15 |
| Isoform 2C2A of Collagen alpha-2(VI) chain | P12110-2 | COL6A2 | 2.01 | 1.05 |
| Collagen alpha-5(VI) chain | A8TX70;A8TX70-2 | COL6A5 | 0.36 | 0.00 |
| Collagen alpha-1(VII) chain | Q02388;Q02388-2 | COL7A1 | 0.10 | 0.04 |
| Collagen alpha-1(VIII) chain | P27658 | COL8A1 | 0.82 | 0.06 |
| Protein disulfide isomerase CRELD1 | Q96HD1;Q96HD1-2 | CRELD1 | 0.40 | 0.06 |
| Cysteine-rich secretory protein LCCL domain-containing 2 | Q9H0B8;Q9H0B8-2;Q9H0B8-3 | CRISPLD2 | 0.41 | 0.03 |
| Isoform 1 of Elastin | P15502;P15502-1;P15502-10;P15502-13;P15502-2;P15502-4;P15502-9 | ELN | 0.02 | 0.02 |
| Isoform 5 of Elastin | P15502-1;P15502-2;P15502-4;P15502-5 | ELN | 5.30 | 1.26 |
| Isoform 11 of Elastin | P15502-11;P15502-8 | ELN | 6.68 | 1.59 |
| Fibrinogen-like protein 1 | Q08830 | FGL1 | 2.09 | 0.19 |
| Growth arrest-specific protein 6 | Q14393;Q14393-1;Q14393-3;Q14393-4;Q14393-5 | GAS6 | 0.07 | 0.01 |
| Gliomedin | Q6ZMI3 | GLDN | 1.01 | 0.19 |
| Hemicentin-2 | Q8NDA2;Q8NDA2-2;Q8NDA2-4 | HMCN2 | 0.05 | 0.01 |
| Isoform 2 of Insulin-like growth factor-binding protein complex acid labile subunit | P35858;P35858-2 | IGFALS | 1.19 | 0.20 |
| Insulin-like growth factor-binding protein 5 | P24593 | IGFBP5 | 0.12 | 0.12 |
| Immunoglobulin superfamily member 10 | Q6WRI0 | IGSF10 | 0.09 | 0.02 |
| Laminin subunit alpha-1 | P25391 | LAMA1 | 0.02 | 0.01 |
| Latent-transforming growth factor beta-binding protein 3 | Q9NS15;Q9NS15-2 | LTBP3 | 0.07 | 0.01 |
| Isoform 2 of Latent-transforming growth factor beta-binding protein 4 | Q8N2S1;Q8N2S1-2;Q8N2S1-3 | LTBP4 | 0.48 | 0.05 |
| Isoform 2 of Matrilin-2 | O00339;O00339-2;O00339-3;O00339-4 | MATN2 | 0.68 | 0.11 |
| Isoform 2 of Matrix Gla protein | P08493;P08493-2 | MGP | 40.52 | 5.19 |
| Protein NDNF | Q8TB73 | NDNF | 0.34 | 0.21 |
| Isoform 6 of Nephronectin | Q6UXI9-6 | NPNT | 0.14 | 0.02 |
| Netrin-4 | Q9HB63;Q9HB63-2;Q9HB63-3 | NTN4 | 0.37 | 0.02 |
| Papilin | O95428;O95428-2;O95428-4;O95428-5;O95428-6 | PAPLN | 0.39 | 0.04 |
| Proteoglycan 3 | Q9Y2Y8 | PRG3 | 1.18 | 0.03 |
| Somatomedin-B and thrombospondin type-1 domain-containing protein | Q8IVN8 | SBSPON | 0.84 | 0.12 |
| Sushi, nidogen and EGF-like domain-containing protein 1 | Q8TER0;Q8TER0-5 | SNED1 | 0.01 | 0.01 |
| Sushi repeat-containing protein SRPX2 | O60687 | SRPX2 | 0.37 | 0.08 |
| Thrombospondin-4 | P35443 | THBS4 | 0.01 | 0.01 |
| Thrombospondin type-1 domain-containing protein 4 | Q6ZMP0;Q6ZMP0-2;Q6ZMP0-3;Q6ZMP0-4 | THSD4 | 0.14 | 0.01 |
| Isoform 3 of Tenascin | P24821;P24821-3 | TNC | 0.34 | 0.02 |
| Isoform 5 of Tenascin | P24821-5 | TNC | 0.05 | 0.01 |
| Isoform 6 of Tenascin | P24821-6 | TNC | 0.10 | 0.01 |
| Tenascin-R | Q92752;Q92752-2 | TNR | 0.17 | 0.02 |
| Usherin | O75445 | USH2A | 0.07 | 0.01 |
| Isoform V1 of Versican core protein | P13611;P13611-2;P13611-3;P13611-4;P13611-5 | VCAN | 0.93 | 0.05 |
| von Willebrand factor A domain-containing protein 2 | Q5GFL6;Q5GFL6-2;Q5GFL6-3 | VWA2 | 0.11 | 0.03 |
| Extracellular matrix organizing protein FRAS1 | Q86XX4;Q86XX4-2 | FRAS1 | 0.03 | 0.01 |

**Table S4**. **Summary of Collagen Species detected in Azo Fraction**. 32 unique collagen isoforms (UniProt ID) were detected in the Azo fraction.

| **UniProt ID** | **Genes** | **Protein Name** | **% Sequence**  **Coverage** | **Avg**  **(pmol/mg)** | **SE** |
| --- | --- | --- | --- | --- | --- |
| P02452 | COL1A1 | Collagen alpha-1(I) chain | 26.6 | 36.27 | 5.4 |
| P08123 | COL1A2 | Collagen alpha-2(I) chain | 22.4 | 36.17 | 6.5 |
| Q96P44;Q96P44-2;Q96P44-3 | COL21A1 | Collagen alpha-1(XXI) chain | 30.0 | 4.30 | 0.6 |
| Q2UY09 | COL28A1 | Collagen alpha-1(XXVIII) chain | 26.1 | 0.36 | 0.0 |
| P02458;P02458-1 | COL2A1 | Collagen alpha-1(II) chain | 28.3 | 0.04 | 0.0 |
| P02461;P02461-2 | COL3A1 | Collagen alpha-1(III) chain | 25.4 | 2.00 | 0.2 |
| P02462 | COL4A1 | Collagen alpha-1(IV) chain | 31.6 | 17.96 | 0.5 |
| P08572 | COL4A2 | Collagen alpha-2(IV) chain | 24.5 | 9.63 | 1.2 |
| Q01955 | COL4A3 | Collagen alpha-3(IV) chain | 27.9 | 2.30 | 1.0 |
| P53420 | COL4A4 | Collagen alpha-4(IV) chain | 26.9 | 1.45 | 0.1 |
| P29400;P29400-2 | COL4A5 | Collagen alpha-5(IV) chain | 31.9 | 0.27 | 0.0 |
| Q14031;Q14031-2 | COL4A6 | Collagen alpha-6(IV) chain | 25.6 | 0.02 | 0.0 |
| P20908;P20908-2 | COL5A1 | Collagen alpha-1(V) chain | 27.7 | 0.44 | 0.0 |
| P05997 | COL5A2 | Collagen alpha-2(V) chain | 27.3 | 0.60 | 0.0 |
| P25940 | COL5A3 | Collagen alpha-3(V) chain | 26.8 | 1.23 | 0.2 |
| P12109 | COL6A1 | Collagen alpha-1(VI) chain | 19.0 | 54.97 | 3.6 |
| P12110;P12110-2;P12110-3 | COL6A2 | Collagen alpha-2(VI) chain | 20.2 | 53.11 | 4.7 |
| P12110-2 | COL6A2 | Isoform 2C2A of Collagen alpha-2(VI) chain | 20.2 | 1.80 | 0.9 |
| P12111;P12111-2;P12111-3;P12111-4;P12111-5 | COL6A3 | Collagen alpha-3(VI) chain | 19.6 | 16.80 | 1.9 |
| P12111-4 | COL6A3 | Isoform 4 of Collagen alpha-3(VI) chain | 18.4 | 1.26 | 0.2 |
| A8TX70;A8TX70-2 | COL6A5 | Collagen alpha-5(VI) chain | 17.5 | 0.36 | 0.0 |
| A6NMZ7;A6NMZ7-2 | COL6A6 | Collagen alpha-6(VI) chain | 15.2 | 3.43 | 0.4 |
| Q02388;Q02388-2 | COL7A1 | Collagen alpha-1(VII) chain | 22.9 | 0.10 | 0.0 |
| P27658 | COL8A1 | Collagen alpha-1(VIII) chain | 34.0 | 0.82 | 0.1 |
| Q03692 | COL10A1 | Collagen alpha-1(X) chain | 27.9 | 0.30 | 0.0 |
| P12107;P12107-2;P12107-3;P12107-4 | COL11A1 | Collagen alpha-1(XI) chain | 27.2 | 0.08 | 0.0 |
| Q99715;Q99715-2;Q99715-4 | COL12A1 | Collagen alpha-1(XII) chain | 19.2 | 1.98 | 0.2 |
| Q05707;Q05707-2;Q05707-3 | COL14A1 | Collagen alpha-1(XIV) chain | 18.8 | 6.51 | 1.0 |
| P39059 | COL15A1 | Collagen alpha-1(XV) chain | 20.7 | 1.55 | 0.1 |
| Q07092;Q07092-2 | COL16A1 | Collagen alpha-1(XVI) chain | 28.8 | 0.19 | 0.0 |
| P39060;P39060-1;P39060-2 | COL18A1 | Collagen alpha-1(XVIII) chain | 26.9 | 3.56 | 0.2 |

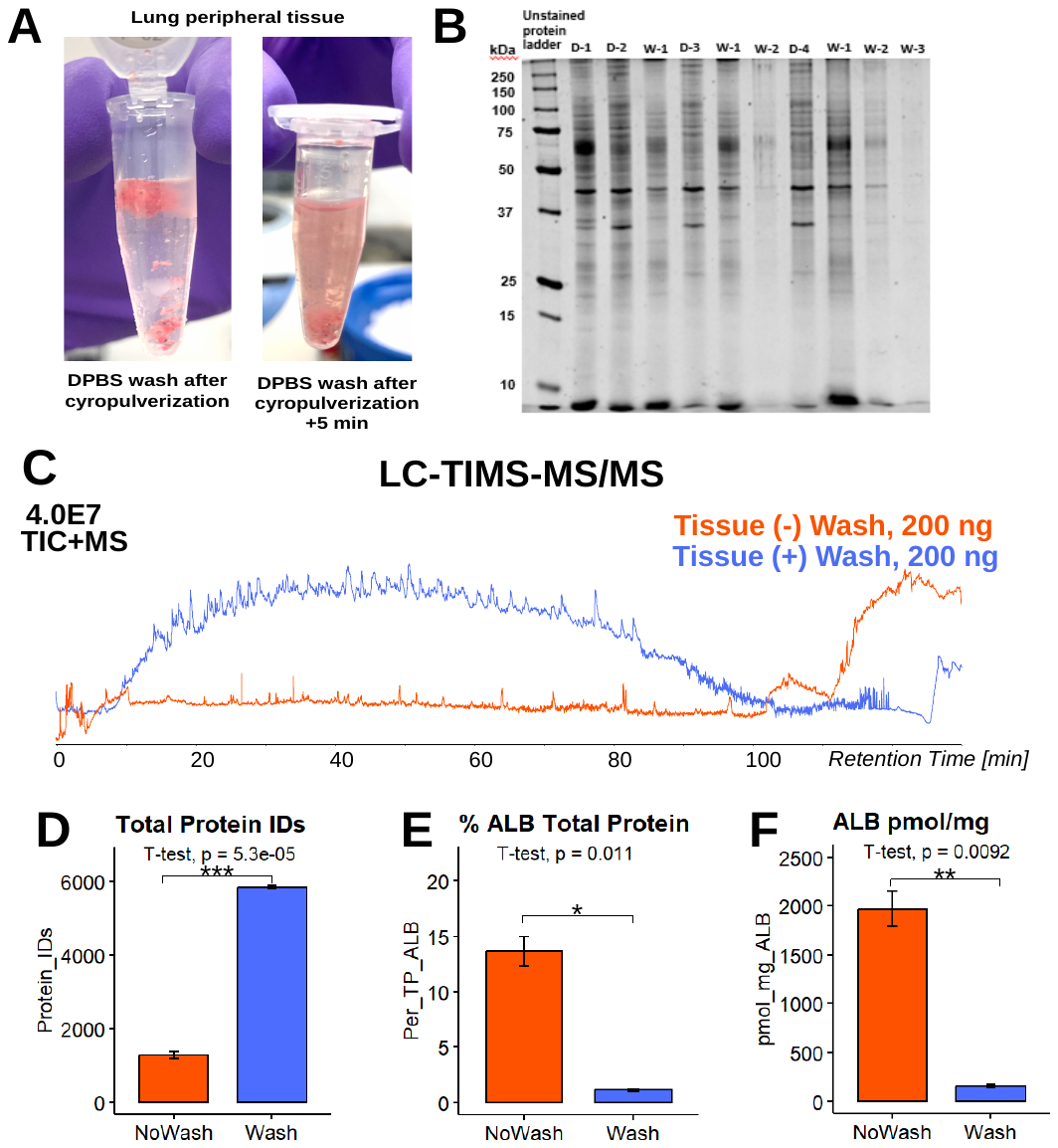

**Figure S1.** **DPBS Wash Post-Cryopulverization Protein Increases Total Protein Signal by Depleting Plasma Proteins. A)** Cryopulverized lung tissue photographed immediately upon submersion in 1 mL DPBS (right) and after 5 min of incubation with DPBS (left). After 5 min, the tissue is gently pelleted by centrifugation and the DPBS is removed. **B)** SDS-PAGE (12.5% SDS, 2 μg total protein loaded each lane, Sypro Ruby stained) of DPBS washes (“W”) and the resulting decellularized protein extracts (“D”). Unwashed lung tissue extract (“D-1”) shows highly abundant protein close to ~70 kDa, which was assigned as Human Serum Albumin (69.4 kDa, UniProt ID: P02768, ALBU_HUMAN). DPBS washes (“W”) and the resulting decellularized protein extracts (“D”) are shown in lanes 3-11, representing one, two, and three sequential washings of lung tissue. **C)** Total Ion Chromatograms (TICs) of LC-MS/MS Decell extracts from the same lung tissue tested with no wash (orange trace) and after washing with DPBS (blue trace). 200 ng peptides were injected to the column for each run. **D)** Comparison of protein identifications in Decell samples (n=3 each group) created with unwashed lung tissue (“No Wash,” orange) and lung tissue washed with DPBS (“Wash,” blue). An average of 1,281 ± 96 proteins were identified in the “No Wash” samples and 5,855 ± 43 proteins were identified in the Washed samples (Mean ± Standard Error, SE, reported). A Student t test was used to determine compare means between groups, and the resulting *p* value is reported. Levels of statistical significance are notated with an asterisk (*): **p* ≤ 0.05, ***p* < 0.01, and ****p* < 0.001; no statistical significance *(ns)* if *p* > 0.05. **E)** Comparison of relative protein intensity of Human Serum Albumin (gene: *ALB*, UniProt ID P02768) intensity in Decell LC-MS/MS samples prepared from tissues that did not undergo a wash (“No Wash,” orange) and those that did undergo a DPBS wash protocol (“Wash,” blue). Relative protein intensity was calculated using the “Total Protein” approach,^1, 2^ and is expressed as a percentage of the total protein intensity. The average % of total protein signal in “No Wash” was 14% ± 1.4 total signal, and ALB comprised 1.1% ± 0.1 total signal in “Wash.” Level of significance between conditions was evaluated by Student t test, and the resulting *p* value is reported. **F)** Comparison of ALB intensity expressed as pmol/g total protein, calculated using the “Total Protein” Approach. Level of significance between conditions was evaluated by Student t test, and the resulting *p* value is reported.

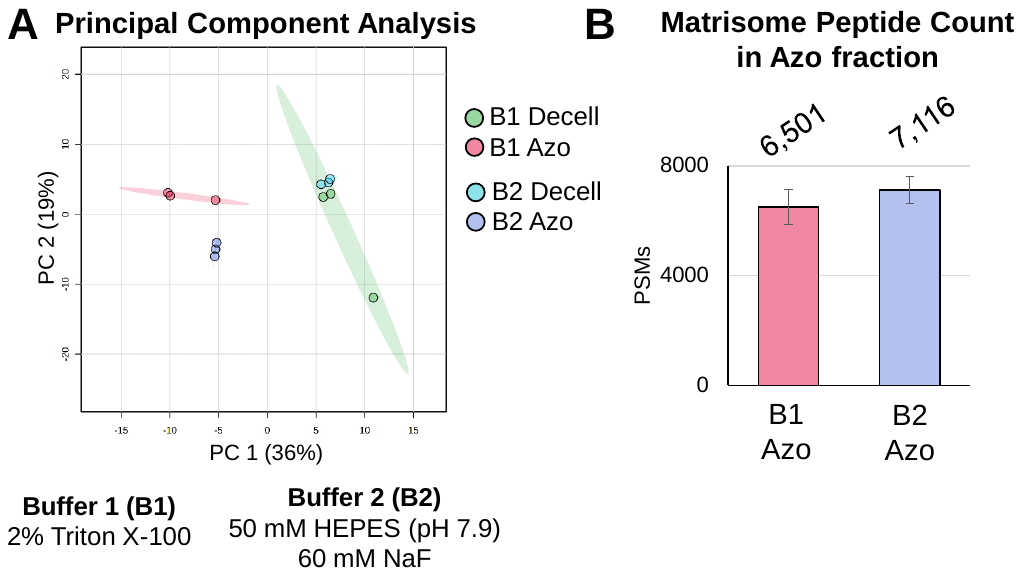

**Figure S2**. **Comparison of HEPES and Triton X-100 based Decellularization of Lung Tissue A)** Principle Component Analysis of LC-MS/MS results. **B)** Comparison of Matrisome Peptide Spectral Matches (PSMs) identified in the Azo extract following a Triton X-100 decellularization (pink) and HEPES-based decellluarization (blue). On average, 6,501 matrisome PSMs were identified in the Triton preparation, and 7,116 matrisome PSMs were identified in the preparation following HEPES-based decellularization.

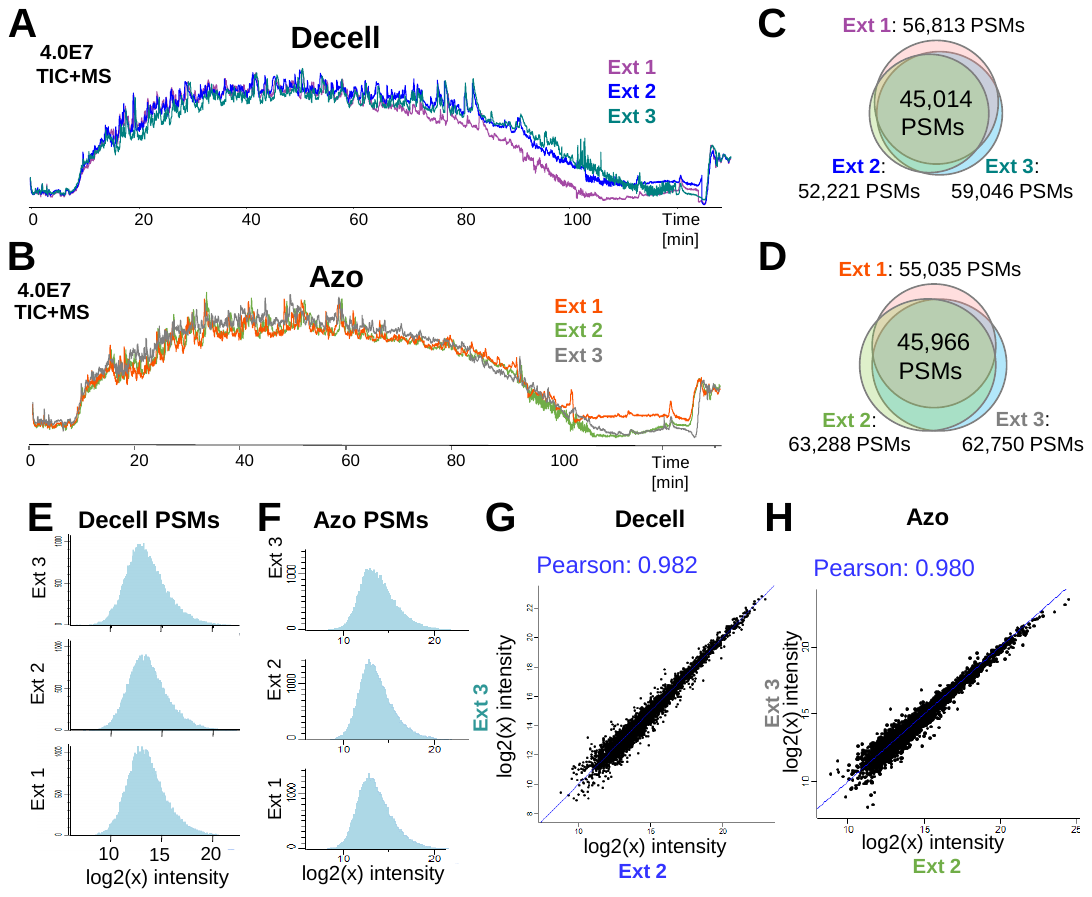

**Figure S3. Strong Reproducibility between Extraction Replicates Observed for all Identified Proteins. A)** Stacked overlay of TIC+MS signal for (n=3) Extraction Replicates of Decell fractions and **B)** corresponding Azo fractions from peripheral lung tissue. 200 ng was peptide injected per run. **C)** Number of Peptide Spectral Matches (PSMs) in each extraction replicate in the Decell fraction. PSMs ranging from 52,221-59,046, with an overlap of 45,014 PSMs identified in every replicate. **D)** Overlap of PSMs identified in each extraction replicate in the Azo fraction. PSMs ranging from 55,035-63,288, with an overlap of 45,966 PSMs identified in every replicate. E) Normal, unimodal distribution of log2x-transformed peptide intensities was observed in extraction replicates for the Decell fractions and **E)** corresponding Azo fractions. **G)** Strong correlation of log2x-transformed protein intensity for all identified proteins in each extraction replicate demonstrates strong extraction reproducibility. Pearson correlation coefficient ranged from 0.965-0.982 for each comparison in the Decell fractions and **H)** 0.960-0.980 in the Azo fractions.

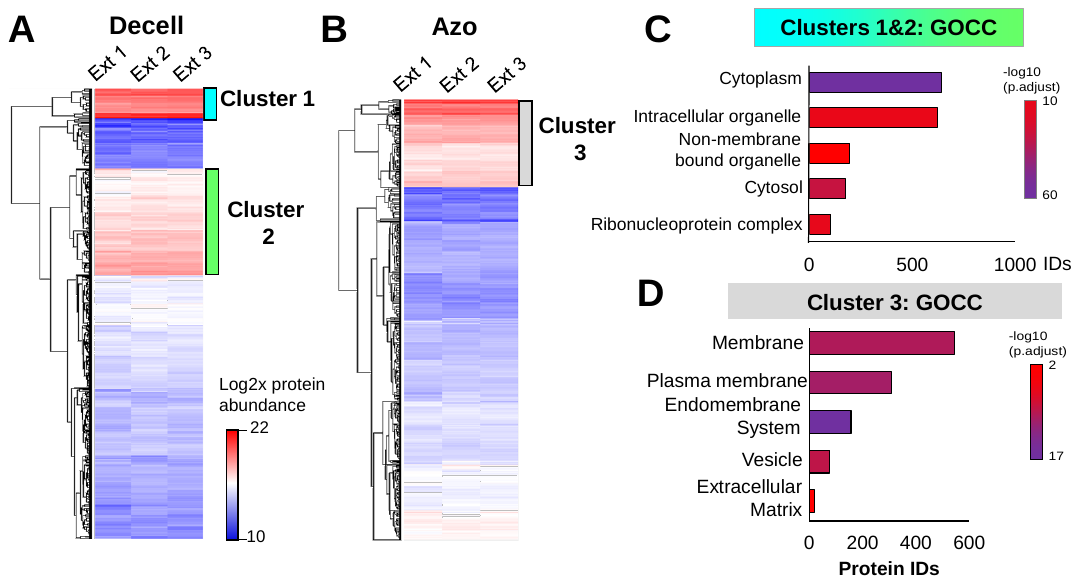

**Figure S4. Global Overview of Decell and Azo Fraction Content. A)** Hierarchical clustering of total proteins identified in the Decell fraction across n=3 extraction replicates. Log2(x)-transformed protein intensity is shown; clusters 1 and 2 represent the highest abundance proteins. **B)** Hierarchical clustering of total proteins identified in the Azo fraction across n=3 extraction replicates. Cluster 3 in the Azo fraction represent the highest abundance proteins. **C)** Gene ontology – cellular compartment overrepresentation analysis of Clusters 1 and 2 highlighted in panel E (1,624 proteins). GOCC terms in this cluster related to cytosolic proteins and organelles, such as ribosomes. **D)** GOCC overrepresentation analysis of Cluster 3, the highest abundance proteins in the Azo fraction. GOCC terms in this cluster related to cytosolic proteins and organelles, such as ribosomes.

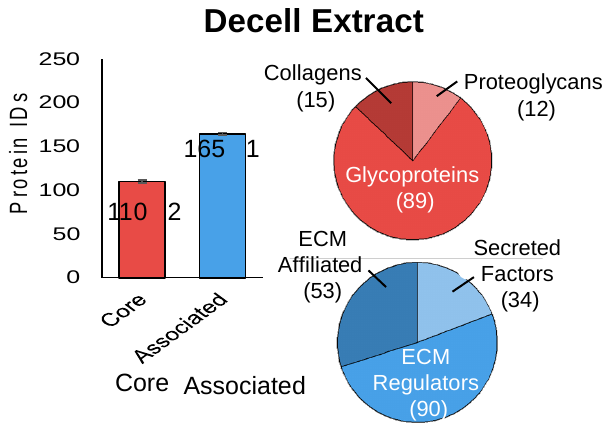

**Figure S5. Overview of Matrisome Recovery in the Decell Fraction.** An average of 110 ±2 core matrisome proteins and 165 ±1 Matrisome-associated proteins were identified between n=3 extraction replicates of peripheral tissue. The core matrisome species consisted of 15 collagens, 12 proteoglycans, and 89 glycoproteins. The matrisome associated proteins consisted of 53 ECM-affilated proteins, 34 secreted factors, and 90 ECM Regulators.

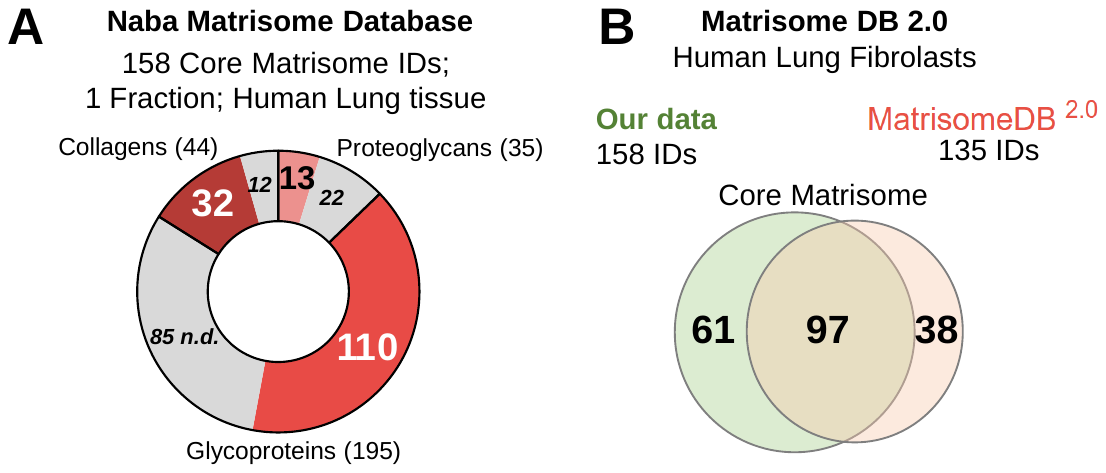

**Figure S6.** **Comparison of core matrisome identifications by Azo the predicted and experimentally observed matrisome.** **A)** Core matrisome identifications made in the Azo fraction (in shades of red) compared to the *in silico* Matrisome Database.^3^ The total number of predicted core matrisome genes for collagens, glycoproteins, and proteoglycans are 44, 35, and 195 IDs, respectively. B) Comparison of core matrisome identifications to experimentally observed core matrisome proteins from the updated Matrisome DB.^4^ One literature report matched to “human lung;” the data is sourced from Lansky, Z. *et. al*. JSBX. **2019**.^5^

**References**

(1) Wiśniewski, J. R.; Rakus, D. Multi-enzyme digestion FASP and the ‘Total Protein Approach’-based absolute quantification of the Escherichia coli proteome. *Journal of Proteomics* **2014**, *109*, 322-331. DOI: <https://doi.org/10.1016/j.jprot.2014.07.012>.

(2) Wiśniewski, J. R. Chapter Four - Label-Free and Standard-Free Absolute Quantitative Proteomics Using the “Total Protein” and “Proteomic Ruler” Approaches. In *Methods in Enzymology*, Shukla, A. K. Ed.; Vol. 585; Academic Press, 2017; pp 49-60.

(3) Naba, A.; Clauser, K. R.; Hoersch, S.; Liu, H.; Carr, S. A.; Hynes, R. O. The Matrisome: <em>In Silico</em> Definition and <em>In Vivo</em> Characterization by Proteomics of Normal and Tumor Extracellular Matrices *<sup> &#x9;&#x9;</sup>. *Molecular & Cellular Proteomics* **2012**, *11* (4). DOI: 10.1074/mcp.M111.014647 (acccessed 2023/06/18).

(4) Shao, X.; Gomez, C. D.; Kapoor, N.; Considine, J. M.; Grams, C.; Gao, Y.; Naba, A. MatrisomeDB 2.0: 2023 updates to the ECM-protein knowledge database. *Nucleic Acids Research* **2023**, *51* (D1), D1519-D1530. DOI: 10.1093/nar/gkac1009 (acccessed 3/22/2023).

(5) Lansky, Z.; Mutsafi, Y.; Houben, L.; Ilani, T.; Armony, G.; Wolf, S. G.; Fass, D. 3D mapping of native extracellular matrix reveals cellular responses to the microenvironment. *Journal of Structural Biology: X* **2019**, *1*, 100002. DOI: <https://doi.org/10.1016/j.yjsbx.2018.100002>.
